# Automatic deep learning driven label-free image guided patch clamp system for human and rodent in vitro slice physiology

**DOI:** 10.1101/2020.05.05.078162

**Authors:** Krisztian Koos, Gáspár Oláh, Tamas Balassa, Norbert Mihut, Márton Rózsa, Attila Ozsvár, Ervin Tasnadi, Pál Barzó, Nóra Faragó, László Puskás, Gábor Molnár, József Molnár, Gábor Tamás, Peter Horvath

## Abstract

Patch clamp recording of neurons is a labor-intensive and time-consuming procedure. We have developed a tool that fully automatically performs electrophysiological recordings in label-free tissue slices. The automation covers the detection of cells in label-free images, calibration of the micropipette movement, approach to the cell with the pipette, formation of the whole-cell configuration, and recording. The cell detection is based on deep learning. The model was trained on a new image database of neurons in unlabeled brain tissue slices. The pipette tip detection and approaching phase use image analysis techniques for precise movements. High-quality measurements were performed on hundreds of human and rodent neurons. We also demonstrate that further molecular and anatomical analysis can be performed on the recorded cells. The software has a diary module that automatically logs patch clamp events. Our tool can multiply the number of daily measurements to help brain research.

**ONE SENTENCE SUMMARY:** Novel deep learning and image analysis algorithms for automated patch clamp systems to reliably measure neurons in human and rodent brain slices.

## INTRODUCTION

Research of the past decade uncovered the unprecedented cellular heterogeneity of the mammalian brain. It is well accepted now, that the complexity of the rodent and human cortex can be best resolved by classifying individual neurons into subsets by their cellular phenotypes *(1–3)*. By characterizing molecular, morphological, connectional, physiological and functional properties several neuronal subtypes have been defined *(4, 5)*. Revealing cell-type heterogeneity is still incomplete and challenging since classification based on quantitative features requires large amount of individual cell samples, often thousands or more, encompassing a highly heterogeneous cell population. Recording morphological, electrophysiological, and transcriptional properties of neurons requires different techniques combined on the same sample such as patch clamp electrophysiology, post hoc morphological reconstruction or single-cell transcriptomics. The fundamental technique to achieve such trimodal characterization of neurons is the patch clamp recording, which is highly laborious and expertise intense. The quantitative and qualitative efficiency of single-cell patch clamp procedure is highly determinant for every follow up measurements including anatomical reconstruction and molecular analyses. Therefore, there is a high demand to efficiently automate this labour intense and challenging process.

Recently, patch clamp technique has been automated and improved to a more advanced level *(6, 7)*. ‘Blind patch clamping’ moves the pipette forward *in vivo (8–10)* and the brain cells are automatically detected by a resistance increase at the pipette tip. Automated systems soon incorporated image-guidance by using multi-photon microscopy on genetically modified rodents *(11–13)*. Further improvements include the integration of tools for monitoring animal behavior *(14)*, the design of an obstacle avoidance algorithm before reaching the target cell or the development of a pipette cleaning method which allows the immediate reuse of the pipettes up to ten times *(16, 17)*. Automated multi-pipette systems were developed to study the synaptic connections *(18, 19)*. It is also shown that cell morphology can be examined using automated systems *(20)*. One crucial step for image-guided automation is pipette tip localization. Different pipette detection algorithms were compared previously *(21)*. The other crucial step is the automatic detection of the cells which has only been performed in two-photon images so far. It is currently not possible to efficiently fluorescently stain human brain tissues. Alternatively, detection of cells in label-free images would open up new application possibilities *in vitro (22)*, e.g. experiments on surgically removed human tissues. Most recently, deep learning*(23)* has been emerging to a level that in the case of well-defined tasks, outperforms humans, and often reaches human performance on ill-defined problems like detecting astrocyte cells *(24)*.

In this paper we describe a system we developed in order to overcome time-consuming and expertise-intense neuron characterization and collection. This fully automated differential interference contrast microscopy (DIC, or label-free in general) image-guided patch clamping system (DIGAP) combines 3D infrared video microscopy, cell detection using deep convolutional neural networks and a glass microelectrode guiding system to approach, attach, break-in and record biophysical properties of the target cell.

The steps of the visual patch clamp recording process are illustrated in Fig. 1a. Before the first use of the system, the pipette has to be calibrated, so that it can be moved relative to the field of view of the camera (1). Thereafter, a position update is made after every pipette replacement (2) using the built-in pipette detection algorithms (3) to overcome the problem caused by pipette length differences. At this point, the system is ready to perform patch clamp recordings. We have acquired and annotated a single cell image database on label-free neocortical brain tissues, to our knowledge the largest 3D set of this kind. A deep convolutional neural network was trained for cell detection. The system can automatically select a detected cell for recording (4). When a cell is selected, multiple subsystems are started simultaneously that perform the patch clamping:

i. A subsystem is controlling the movement of the micropipette next to the cell. If any obstacle is found in the way, an avoidance algorithm tries to bypass it (5).
ii. A cell tracking system follows the possible shift of the cell in 3D (7).
iii. During the whole process, a pressure regulator system assures that the requested pressure on the pipette tip is available (6).

**Figure 1.**
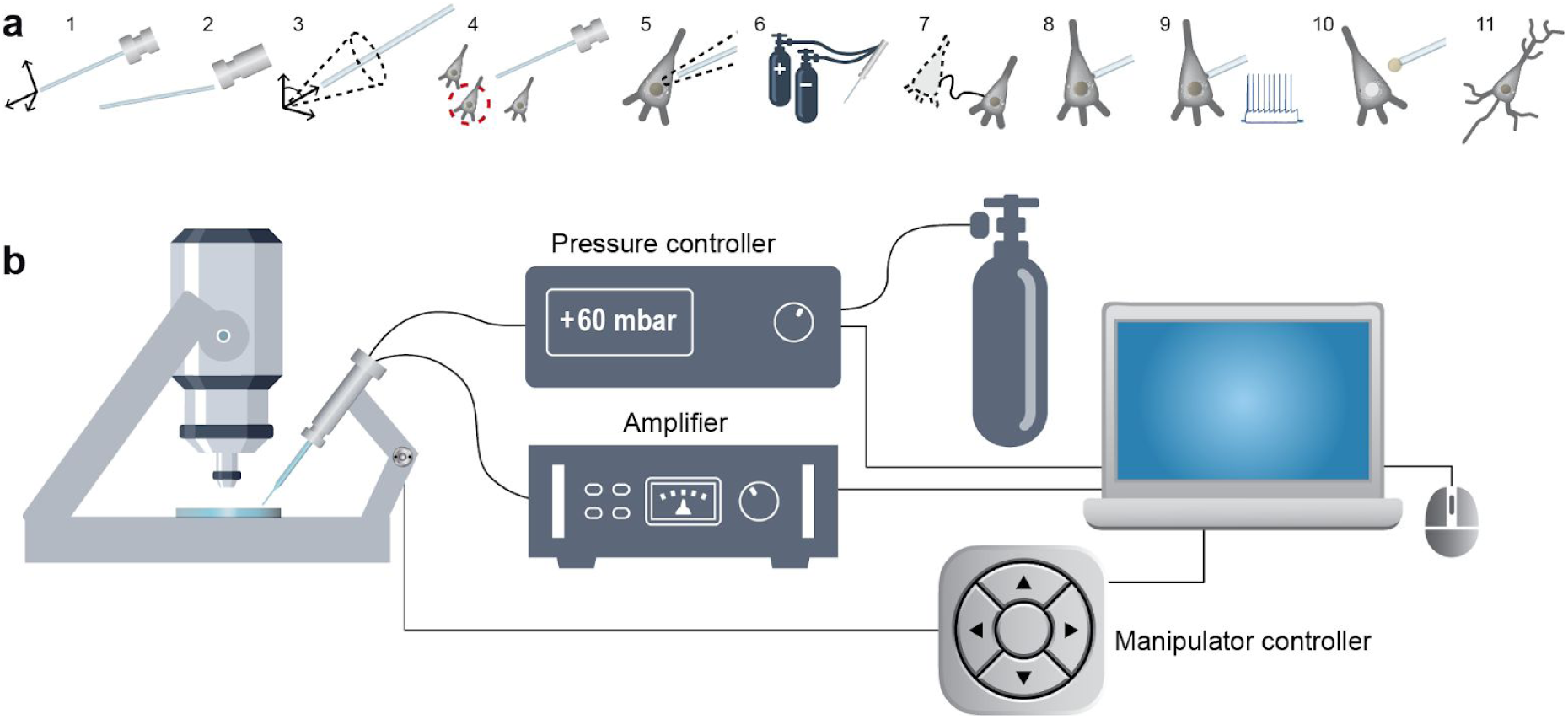
Scheme of the DIC image-guided automatic patch clamp (DIGAP) system. **a:** Steps of DIGAP procedures. 1: Pipette calibration by the user, 2: pipette replacement after recording, 3: image-based automatic pipette tip detection, 4: automatic cell detection, 5: pipette navigation to the target cell, 6: pressure regulation, 7: 3D cell tracking, 8: gigaseal formation, 9: break-in and electrophysiological recording, 10: nucleus and cytoplasm harvesting, 11: anatomical reconstruction of the recorded cell. **b:** Hardware setup of the DIGAP system.

Once the pipette touches the cell (cell-attached configuration) the system performs gigaseal formation (8), then breaks in the cell membrane and automatically starts the electrophysiological measurements (9). The nucleus or the cytoplasm of the patched cell can be harvested (10). Later, the recorded cells can be anatomically reconstructed in the tissue (11).

At the end of the measurements, the implemented pipette cleaning method can be performed or the next patch clamp recording can be started after pipette replacement and from the pipette tip position update step (3). An event logging system collects information during the patch clamp process, including the target locations and the outcome success, and report files can be generated at the end. The report files are compatible with the Allen Cell Types Database *(25)*. Our system was tested on rodent and human samples *in vitro*. The quality of the electrophysiological measurements strongly correlates to that of made by a trained experimenter. We used the system for harvesting cytoplasm and nucleus from the recorded cells and performed anatomical reconstruction on the samples. Our system is the first that can operate on unstained tissues using deep learning, that reaches and even outperforms the cell detection accuracy of human experts, and that enables the multiplication of the number of recordings while preserving high-quality measurements.

## RESULTS

Here we introduce an automated seek-and-patch system that performs electrophysiological recordings and sample harvesting for molecular biological analysis from single cells on unlabeled neocortical brain slices. Using deep learning, trained on a previously built database of single neurons acquired in 3D, our system can detect most of the healthy neuronal somata in a Z-stack recorded by DIC microscopy from a living neocortical slice. The pipette approaches the target cell, touches it, acquires electrophysiological data, and the cell’s nucleus can be isolated for further molecular analysis. Components of the system are a typical electrophysiological setup: IR video microscopy imaging system, motorized microelectrode manipulators, XY shifting table, electrical amplifier and a custom-designed pressure controller. All these elements were controlled by a custom-developed software (available at https://bitbucket.org/biomag/autopatcher/). The system was successfully applied to perform patch clamp recordings on a large set of rodent and human cells (100 and 74 respectively). The automatically collected cells well represent the wide-range phenotypic heterogeneity of the brain cortex. Subsequent transcriptome profiling and whole-cell anatomical reconstruction confirmed the usefulness and applicability of the proposed system.

In this section, we present the hardware components, the developed computational methodologies and the biological evaluation of our system.

### Hardware development and control

The schematic hardware setup of the proposed system is shown in Fig. 1b. The software system we developed controls each hardware using their drivers on application programming interface (API) level, which makes the system modular and different types of hardware component (e.g. manipulators, biological amplifier, XZ shifting table etc.) can be attached. The electrophysiological signal is transferred to the DIGAP software via USB digitizer board (National Instruments, USB-6009). To apply different air pressure on the pipette in distinct phases of the patching procedure we built a custom pressure controller detailed in Supplementary Information: Pressure Regulator. Analog pressure sensors are used for monitoring the actual air pressure on the pipette and voltage signals of the sensors were connected in the input channels of the USB digitizer board. The solenoid valves of the regulator are controlled with TTL signals of the digital output channels of the digitizer.

### Pipette calibration and automatic detection

Pipette calibration is a one-time process which determines the coordinate system transformation between the pipette and the stage axes. The calibration consists of moving the pipette along its axes with known distances, finding it with the stage and detecting the exact pipette tip position in the camera image. Calibration allows the pipette to be moved at any position of the microscope stage space. Note that no assumptions are made on the orientation or the tilt angles of the pipette.

The glass pipettes usually differ in length, thus the tip position should be updated after a pipette change. To automate this step we have developed algorithms for pipette detection in DIC images. First, we use a fast initialization heuristic and then refine the detection. The refinement step is the extension of our previous differential geometry-based method to 3 dimensions *(21)*. The pipette is modeled as two cylinders that have a common reference point and an orientation. The model is updated by the gradient descent method such that it covers dark regions introduced by the pipette in the image. Fig. 2a shows the starting and final state of the algorithm from different projections in gradient images for visualization purposes. The detailed description of the algorithms and the equation derivations can be found in Supplementary Information: Pipette Detection System. The algorithm has an accuracy of 0.99 ± 0.55 μm compared to manually selected tip positions, which allow to reliably reach the cells of 10 μm diameter (on average) with the pipette when oriented towards their centroids.

**Figure 2.**
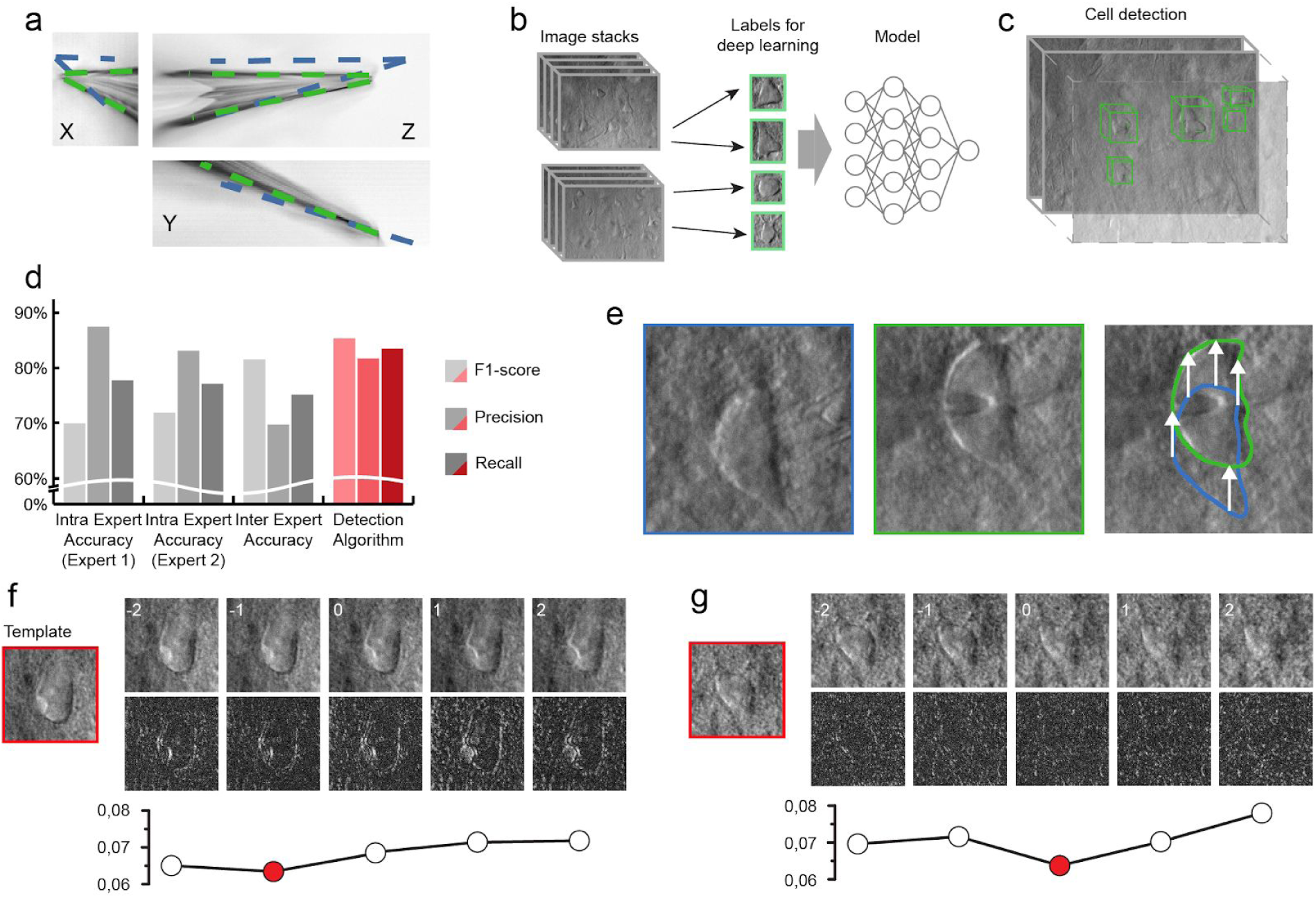
The developed algorithms for the DIGAP system. **a:** Result of the Pipette Hunter detection model shown in three different projections of the image stack. Blue lines show the initial state of our pipette localization algorithm, while green lines show the result of the method. **b:** Training dataset generation: 265 image stacks (60-100 images per stack with 1 μm frame distance along the Z axis) captured from human and rodent neocortical slices with DIC videomicroscopy (left). 31,720 objects as healthy cells (green boxes) labeled on every slice of the image stack by 4 experts. **c:** After training session the DIGAP system detects cells in unstained living neocortical tissues. **d:** Accuracy of the automated cell detection pipeline. Both intra- and inter expert measurements are performed. **e:** Lateral tracking of the cell movement. DIC images of the targeted (in blue box) and patched cell (in green box). The cell drifted from its initial location (yellow arrows in the right panel) during the pipette maneuver. **f-g:** Z-tracking of the cell movement. The template image was captured at the optimal focal depth (in red boxes) before starting the tracking. During the pipette movement image stacks were captured from the targeted cell (upper panels). The bottom row shows the differences between the template and the image of the corresponding z position. The lowest standard deviation value of the difference images (plots) shows the direction of the cell drift in the Z axis.

### Cell detection

We applied a deep learning algorithm in order to detect cells in DIC images and propose them for automatic patch clamp recording. Object detection of neurons in label-free tissue images is challenging *(24)*. Various software solutions *(26, 27)* were developed to segment neurons in cell cultures, however, they do not provide satisfactory results on tissues. To obtain a reliable object detection in brain tissue, we designed a cell detection algorithm, which involved three steps: data annotation, training of the model and inference.

The annotation of the image stacks was performed by 4 field experts using our custom-built annotation software, which resulted in 31,720 objects labeled as healthy cells. The training process was capable of generating a suitable model that recognizes neurons in their original environment in DIC images (Fig. 2b). Our algorithm detects neurons in 2D images, then it extends the detection along the z-axis in the image stacks to complete the object detection in 3D space (Fig. 2c). Four different comparisons were performed for evaluation. Intra-expert accuracies were measured by showing a small part of the dataset to the annotators again after 3 months. Inter-expert accuracy was also measured to compare the annotators and resulted in a 0.752 F1-score. The deep learning model outperformed both annotators with a 0.835 F1-score (81.72% precision and 85.39% recall, Fig. 2d).

The result model was imported to the software. When the user initiates cell detection a stack is created and the detected cells are highlighted with bounding boxes (Fig 2c). The detections are ordered by the confidence value thus healthier cells are offered earlier. The target cell can also be selected manually based on arbitrary criteria required for the experiment. The details of the detection system can be found in Supplementary Information: Cell Detection System.

### Tracking the cell in 3D

Due to the elasticity of the tissue the movement of the pipette can significantly deform it and change the location of the cell of interest. In order to precisely re-define the pipette trajectory, the location of the target cell needs to be tracked. We have developed an online system that performs tracking in the lateral and Z directions (Fig. 2e-g). Both directions require a template image of the target cell which is acquired before starting the patch clamp process when the cell is in the focal plane of the microscope. The lateral tracking is performed in the image of the most recent focal level. It uses the Kanade-Lucas-Tomasi (KLT) feature tracker algorithm *(28, 29)*. The Z tracking is based on a focus detection algorithm that operates on a small image stack encompassing target cell body. The standard deviation of the images of the target cell body is computed and compared to initial images. As a result, the displacement direction of the target cell along the Z axis is determined. The whole process was done with stopped pipette to ensure that the cell is not pushed away meanwhile. The detailed explanation of the algorithms with examples can be found in Supplementary Information: Cell Tracking System.

### Automated patch clamping steps

After pipette calibration and cell detection the patch clamping procedure can be started. First, the DIGAP software calculates the trajectory of the pipette movement along which the manipulator moves the pipette tip close to the cell while applying medium air pressure (50-70 mbar). The initial trajectory is a straight line along the manipulators X axis. Note that the X axis of the manipulator is tilted so the movement vector of the pipette is parallel to the longitudinal axis of the pipette. We found that approaching is more reliable if the pipette is first moved a few micrometers above the cell and then finally descending on it. The impedance of the pipette tip is monitored continuously during the movement.

During the movement of the pipette air pressure is dynamically changed with predefined air pressure values. Air pressures were empirically set for the different phases: hunting, sealing, and breaking. Pipette tip impedance was continuously checked in order to detect phases and apply the task specific pressure.

Early resistance increase denotes the presence of an obstacle in front of the pipette, e.g. a blood vessel or another cell. If an obstacle is hit, the pipette is pulled back, slightly moved laterally and when the obstacle is passed the pipette is oriented back to the initial trajectory towards the target *(15)*. Meanwhile, the described 3D tracking algorithm compensates for the movement trajectory due to the possible displacement of the target cell. When the pipette tip reaches the target position above the cell, the pressure is decreased to a low positive value (10-30 mbar). Then the pipette is moved in Z direction and the resistance of the tip is monitored by 5 ms long -5 mV voltage steps. If the impedance increases more than a predefined value (0.7-1.2 MΩ) the sealing phase is initiated. The cell-attached configuration is set up by the immediate cease of pressure. To achieve tight sealing of the cell membrane into the glass we apply small negative pressure (from -30 to -10 mbar) and the holding potential is set to -60 mV stepwise. If seal resistance reaches 1 GΩ (‘gigaseal’) then suction pulses of increasing length are applied to break-in the membrane. Information about the process, including pipette distance from the target, actual air pressure, and electrical resistance values are continuously monitored and shown in the GUI windows. Description of the steps and the parameter values are described in detail in Supplementary Information: Software Usage. A representative procedure is demonstrated in Fig. 3.

**Figure 3.**
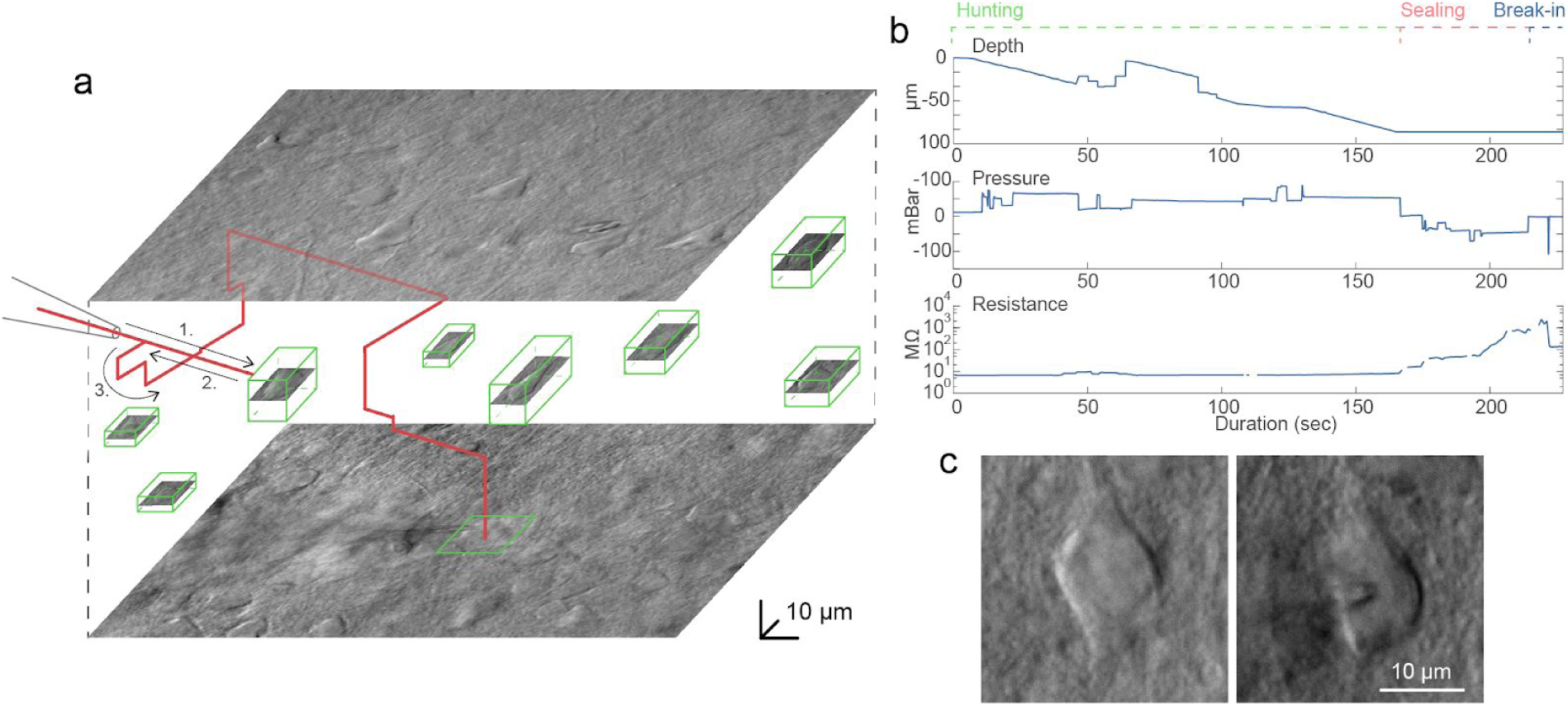
Representative example of a whole-cell recording. **a:** Trajectory of the pipette tip (red line) with obstacle avoidance (numbered) in the tissue and the spatial location of the detected cells (green boxes). **b:** Plots of the depth of the pipette tip in the tissue, the applied air pressure, and the measured pipette tip resistance during the approach. **c:** Image of a cell before and after performing patch clamp recording on it.

### Software

The control software is written in MATLAB and the source code is made publicly available at https://bitbucket.org/biomag/autopatcher/. The visual patch clamping process can be started from a user-friendly GUI which allows every parameter to be set and the process to be monitored real-time by the operator. Throughout the session, the Patch Clamp Diary module collects and visualizes information about patch clamping attempts, including their location and outcome status. The user can additionally mark positions in the biological sample that help orientation during the experiment (i.e. boundaries of the brain slice or the parallel strands that keep secure the tissue).

Many utility features are present to help everyday experimenting. Single images or image stacks can be acquired, saved or loaded from the menu bar. The acquired images can be processed by performing background illumination correction or DIC image reconstruction which can help in identifying cells and their features. The graphical processing unit (GPU) extension of our reconstruction algorithm *(30)* can be used for reconstruction, which results in about 1000x speed increase. The software contains a built-in labeling tool that allows image database generation to train deep learning cell recognition. Furthermore, most recent practices from other automation systems have also been implemented for the *in vivo* usage, including pipette cleaning*(16, 17)* or hit reproducibility check*(31)*. The XML configuration file makes the adaptation easy between different setups and the software can also operate as a general microscope controller. A logging system is used for maintainability purposes.

### Application in brain slices

To test the performance and effectiveness of our system we obtained a series of recordings on slice preparation of rat somatosensory and visual cortices (n=23 animals) and human temporal and association cortices (n=16 patients). Successful automatic whole-cell patch clamp trials without experimenter assistance were achieved in a total number of n=100 and n=74 (rodent visual and somatosensory cortices and human cortex, respectively) out of n=157 and n=198 attempts. The quality of recordings was supervised by measuring series resistance (R_s_) (Fig. 4). We found a wide range of R_s_ values within successful attempts in both species: 34.52 ± 18.99 MΩ in rat and 31.39 ± 16.67 MΩ in human recordings. Trials with R_s_ value exceeding 100 MΩ were noted as unsuccesful attempts. Access resistance in 48.28% of our recordings was under 30 MΩ which we denoted as high quality and used for further analysis. We applied standard stimulation protocol and recorded membrane potential responses to injected currents. Based on the extracted common physiological features and firing patterns we grouped neurons into electrophysiological types (e-types*(32)*) based on criteria established by the Petilla convention *(33)*. There were 8 e-types in automatic patched rat samples: pyramidal cell (pyr), burst adapting (bAD), continuous non-accommodating (cNAC), continuous stuttering (cSTUT), burst stuttering (bSTUT), delayed stuttering (dSTUT), continuous adapting (cAD), delayed non-accommodating (dNAC). From the human samples, 7 e-types were identified. In our automatically-collected dataset, dNAC type was not represented (Fig. 4).

**Figure 4.**
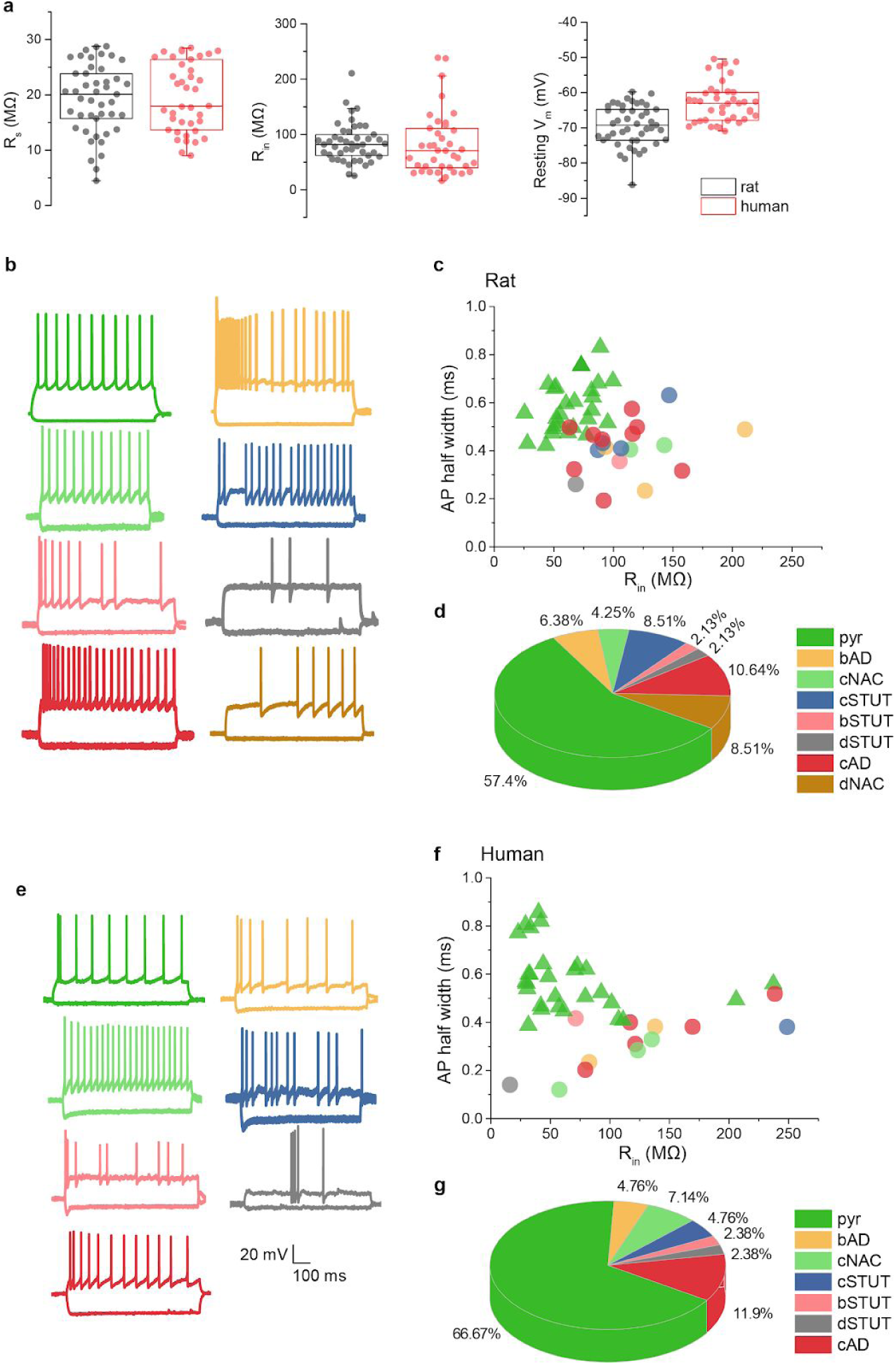
Electrophysiological properties of the cells patched with DIGAP. **a:** Main electrophysiological parameters from the successful automatic patch clamp recordings. The box plots show the series resistance (R_s_,left panel), the input resistance (R_in_, middle panel) and the resting membrane potential (right panel) of all successful measurements. **b:** Different cell types are identified according to firing features: pyramidal cell (pyr), burst adapting (bAD), continuous non-accommodating (cNAC), continuous stuttering (cSTUT), burst stuttering (bSTUT), delayed stuttering (dSTUT), continuous adapting (cAD), delayed non-accomodating (dNAC). **c:** The proportion of recorded cell types. **d:** Individual neurons’ action potential half-widths are presented as a function of the same neuron’s R_in_. Note the segregation of excitatory and inhibitory neuronal classes. Dataset is recorded from rodent samples (Panel c and d colors correspond to panel b). **e-g:** same plots as b-d, representing the dataset recorded in human neocortical slices.

Each electrophysiological recording was performed using biocytin-containing intracellular solution for further anatomical investigation. Among autopatched neurons with <30 MΩ access resistance 80% and 60.71% of full recovery was achieved from rat and human samples, respectively (Fig. 5a).

**Figure 5.**
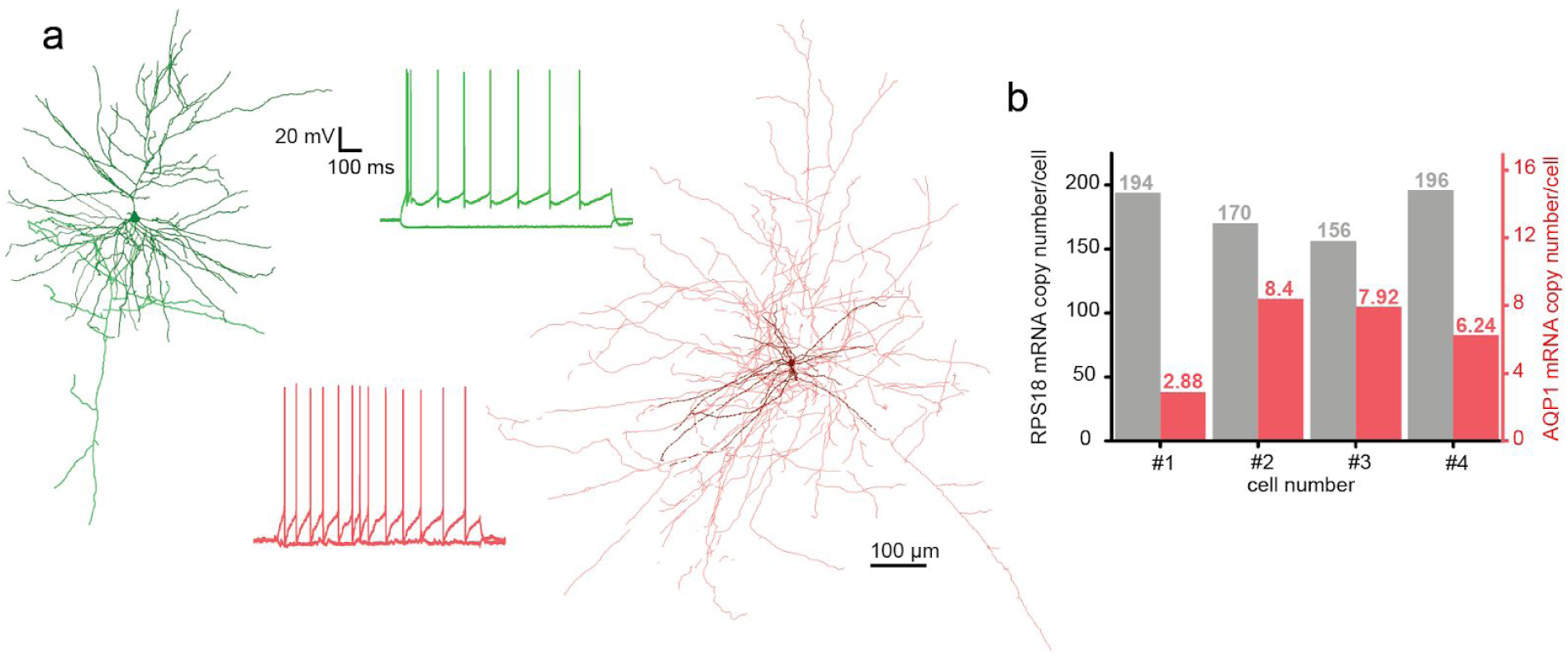
Anatomical and molecular biological investigation of neurons patched with DIGAP. **a:** Two anatomically reconstructed human autopatched neurons. The darker colors represent somata and dendrites of the pyramidal (green) and the interneuron (red) cells. The brighter color shows the axonal arborization. The firing patterns of the cells are the same color as their reconstructions. **b:** mRNA copy numbers of a housekeeping (RPS18, black bars) and the aquaporin 1 (AQP1, red bars) gene from four representative human pyramidal cells.

We next tested if single-cell RNA analysis is achievable from the collected cytoplasm of autopatched neurons. After whole-cell recording of the neurons in the brain slices the intracellular content of the patched cells were aspirated into the recording pipette with gentle vacuum applied by the pressure regulator unit (−40 mBar for 1 min, then -60 mBar for 2-3 min, and finally -40 mBar for 1 min). The tight seal was maintained and the pipette was carefully withdrawn from the cell to form an outside-out configuration. Subsequently, the content of the pipette was expelled into a low-adsorption test tube (Axygen) containing 0.5 μl SingleCellProtectTM (Avidin Ltd. Szeged, Hungary) solution in order to prevent nucleic acid degradation and to be compatible with direct reverse transcription reaction. Then the samples were used for digital polymerase chain reaction (dPCR) analysis to determine the copy number of selected genes. From four single pyramidal cell cytoplasm samples which were extracted from the human temporal cortex, we determined the copy number of a ribosomal housekeeping RPS18 and aquaporin 1 (AQP1) genes (Fig. 5b). The results of the dPCR experiments are in agreement with our previous observations *(34, 35)*.

## METHODS

### Hardware setup

An Olympus BX61 (Tokyo, Japan) microscope with a 40x objective was used for 3D imaging. For moving the pipette and the microscope stage we used Luigs and Neumann Mini manipulators with SM-5 controllers (Ratingen, Germany). The electrophysiological signals were measured by a HEKA EPC-10 amplifier (HEKA Elektronik, Lambrecht, Germany). The signals were digitized at 100 kHz and Bessel filtered at 10 kHz.

### In vitro preparation of human and rat brain slices

All procedures were performed according to the Declaration of Helsinki with the approval of the University of Szeged Ethics Committee. Human slices were derived from materials that had to be removed to gain access for the surgical treatment of deep-brain tumors, epilepsy or hydrocephalus from the association cortical areas with written informed consent of female (n=9, aged 48.2 ± 26.6 years) and male (n=7, aged 48.3 ± 9.9 years) patients prior to surgery. Anesthesia was induced with intravenous midazolam and fentanyl (0.03 mg/kg, 1–2 μg/kg, respectively). A bolus dose of propofol (1–2 mg/kg) was administered intravenously. To facilitate endotracheal intubation, the patient received 0.5 mg/kg rocuronium. After 120 s, the trachea was intubated and the patient was ventilated with a mixture of O_2_ and N_2_O at a ratio of 1:2. Anesthesia was maintained with sevoflurane at monitored anesthesia care (MAC) volume of 1.2–1.5. After surgical removing blocks of tissue were immediately immersed in ice-cold solution containing (in mM) 130 NaCl, 3.5 KCl, 1 NaH_2_PO_4_, 24 NaHCO_3_, 1 CaCl_2_, 3 MgSO_4_, 10 d(+)-glucose, saturated with 95% O_2_ and 5% CO_2_. Slices were cut perpendicular to cortical layers at a thickness of 350 μm with a vibrating blade microtome (Microm HM 650 V) and were incubated at room temperature for 1 hour in the same solution. The artificial cerebrospinal fluid (aCSF) used during recordings was similar to the slicing solution, but it contained 3 mM CaCl and 1.5 mM MgSO_4_.

Coronal slices (350 μm) were prepared from the somatosensory cortex of male Wistar rats (P18-25, n=23, RRID: RGD_2312511)*(36)*. Recordings were performed at 36°C temperature. Micropipettes (3.5–5 MΩ) were filled with low [Cl] intracellular solution for whole-cell patch clamp recording: (in mM) 126 K-gluconate, 4 KCl, 4 ATP-Mg, 0.3 GTP-Na_2_, 10 HEPES, 10 phosphocreatine, and 8 biocytin (pH 7.20; 300 mOsm).

### Molecular biological analysis

After harvesting the cytoplasm of the recorded cells the samples were frozen in dry ice and stored at -80 °C until used for reverse transcription. The reverse transcription (RT) of the harvested cytoplasm was carried out in two steps. The first step took 5 min at 65 °C in a total reaction volume of 5 μl containing 2 μl intracellular solution and SingleCellProtectTM mix with the cytoplasmic contents of the neuron, 0.3 μl TaqMan Assays, 0.3 μl 10 mM dNTPs, 1 μl 5× first-strand buffer, 0.3 μl 0.1 mol/ L DTT, 0.3 μl RNase inhibitor (Life Technologies) and 100 U of reverse transcriptase (Superscript III, Invitrogen). The second step of the reaction was carried out at 55 °C for 1 h and then the reaction was stopped by heating at 75 °C for 15 min. The reverse transcription reaction mix was stored at −20 °C until PCR amplification. For digital PCR analysis the reverse transcription reaction mixture (5 μl), 2 μl TaqMan Assays (Life Technologies), 10 μl OpenArray Digital PCR Master Mix (Life Technologies) and nuclease-free water (5.5 μl) were mixed in a total volume of 20 μl. The mixture was evenly distributed on an OpenArray plate. RT mixes were loaded into 4 wells of a 384-well plate from which the OpenArray autoloader transferred the cDNA master mix by capillary action into 256 nanocapillary holes (4 subarrays) on an OpenArray plate. Processing of the OpenArray slide, cycling in the OpenArray NT cycler and data analysis was done as previously described *(34)*. For our dPCR protocol amplification, reactions with CT confidence values below 100 as well as reactions having CT values less than 23 or greater than 33 were considered primer dimers or background signals, respectively, and were excluded from the data set.

### Anatomical processing and reconstruction of recorded cells

Following electrophysiological recordings, slices were transferred into a fixative solution containing 4% paraformaldehyde, 15% (v/v) saturated picric acid, and 1.25% glutaraldehyde in 0.1 M phosphate buffer (PB; pH=7.4) at 4 °C for at least 12 h. After several washes with 0.1 M PB, slices were frozen in liquid nitrogen then thawed in 0.1 M PB, embedded in 10% gelatin, and further sectioned into 60-μm slices. Sections were incubated in a solution of conjugated avidin-biotin horseradish peroxidase (ABC; 1:100; Vector Labs) in Tris-buffered saline (TBS, pH=7.4) at 4 °C overnight. The enzyme reaction was revealed by 3′ 3-diaminobenzidine tetra-hydrochloride (0.05%) as chromogen and 0.01% H_2_O_2_ as oxidant.

Sections were postfixed with 1% OsO_4_ in 0.1 M PB. After several washes in distilled water, sections were stained in 1% uranyl acetate and dehydrated in an ascending series of ethanol. Sections were infiltrated with epoxy resin (Durcupan) overnight and embedded on glass slides. Three-dimensional light-microscopic reconstructions were carried out using a Neurolucida system (MicroBrightField) with a 100× objective.

### Pipette cleaner

We implemented a pipette cleaning method *(16)* into our system. The cleaning procedure requires two cleaning agents: Alconox, a commercially available cleaning detergent, and artificial cerebrospinal fluid (aCSF). We 3D printed a holder for two PCR tubes containing the liquids that can be attached to the microscope objective and are reachable by the pipette tip. The cleaning is performed by pneumatically taking up and then removing the agents into and from the pipette. The vacuum strength used for the intake of the liquids is -300 mBar and the pressure used for the expulsion is +1000 mBar. The method consists of three steps. First, the pipette is moved to the cleaning agent bath and vacuum is applied for 4 seconds. Then, to physically agitate glass-adhered tissue, pressure and vacuum are alternated, each for 1 second and repeated 5 times total. Finally, pressure is applied for 10 seconds to make sure all detergent is removed. In the second step, the pipette is moved to the aCSF bath and any remaining detergent is expelled by applying pressure for 10 seconds. In the third step, the pipette is moved back to the position near to the biological sample where the cleaning process was initiated. In the original paper, it is shown that these pressure values and the duration of the different steps are more than enough to cycle the volume of agents necessary to clean the pipette tip. We provide a graphical window in our software to calibrate the pipette positions of the tubes containing the cleaning agent and the aCSF and to start the cleaning process.

## DISCUSSION

The developed DIGAP system is able to fully automatically perform whole cell patch clamp recordings on single neurons in rodent and human neocortical slices (Supplementary Video 1). This is a step forward towards characterizing and understanding the phenotypic heterogeneity and cellular diversity of the brain. A central part of the method is the detection of single neurons in label-free 3D images using deep convolutional neural networks reaching super-human precision. The system we developed is fully controlled by a single software, including all hardware components, data handling, and visualization. The control software has its highly comprehensive internal logging system, that allows tracking the approaching and harvesting conditions to such a detailed level. In addition, it can connect to and save database entry records that are compatible with the Allen Brain Atlas single neuron database. In this work, we demonstrated the power of our system that is capable of measuring a large set of rodent and human neurons in the brain cortex. The results show strong correlation to the earlier results in literature in terms of quality and phenotypic composition of cell heterogeneity. Records of measured cells were inserted to the database of the Allen Institute for Brain Science and a subset of the cells was isolated from their tissue environment and single-cell mRNA copy numbers of two selected genes were determined. Furthermore, we successfully demonstrated that autopatched neurons can be anatomically reconstructed.

The main advantage of the proposed system is that it is easily portable to any existing setups and although we do not believe that it will fully substitute human experts, it is a great choice for complex specific tasks, allows parallelization and speeds up discovery. It is important to emphasize the need for a standardized and fully documented patch clamping procedure, which is guaranteed by using DIGAP. The choice of advanced image analysis and deep learning techniques made it possible to work with the least harmful imaging modalities at a human expert level of single-cell detection that was impossible so far. Further possibilities are more widespread and potentially enabling or accelerating discoveries. Combining with intelligent single-cell selection strategies of the detected cells, the proposed system can be the ultimate tool to reveal and describe cellular heterogeneity. In multiple patch clamp setup it can be used to describe the connectome at cellular level. We presented DIGAP’s application to brain research, but other fields, such as cardiovascular or organoid research will benefit from the system. Based on its nearly complete automation, it can help in education.

We plan to enhance our cell detection algorithm with the capability of predicting neuronal phenotypes on label-free DIC images, that will allow targeting specific neuron classes. We also plan to add multi-pipette support to study connections between pairs, triplets or a higher number of cells at a time.

## Supporting information

Supplementary Information: Cell Detection System

Supplementary Information: Cell Tracking System

Supplementary Information: Pipette Detection System

Supplementary Information: Pressure Regulator

Supplementary Information: Software Usage

## ACKNOWLEDGEMENTS

We thank Tímea Tóth and Réka Hollandi for their help in the image labeling, Ádám Szűcs for his work in the early stages of the development, Tamás Szépe for the advice on manipulator control, Nelli Tóth for the anatomical reconstruction, and István Grexa for the 3D printing. This work was supported by NAP-B brain research grant, the NVidia GPU Grant program, the LENDULET-BIOMAG Grant (2018-342) and from the European Regional Development Funds (GINOP-2.3.2-15-2016-00006, GINOP-2.3.2-15-2016-00026, GINOP-2.3.2-15-2016-00037).

## REFERENCES

1. B. Tasic, V. Menon, T. N. Nguyen, T. K. Kim, T. Jarsky, Z. Yao, B. Levi, L. T. Gray, S. A. Sorensen, T. Dolbeare, D. Bertagnolli, J. Goldy, N. Shapovalova, S. Parry, C. Lee, K. Smith, A. Bernard, L. Madisen, S. M. Sunkin, M. Hawrylycz, C. Koch, H. Zeng, Adult mouse cortical cell taxonomy revealed by single cell transcriptomics, Nat. Neurosci. 19, 335–346 (2016).

2. B. Tasic, Z. Yao, L. T. Graybuck, K. A. Smith, T. N. Nguyen, D. Bertagnolli, J. Goldy, E. Garren, M. N. Economo, S. Viswanathan, O. Penn, T. Bakken, V. Menon, J. Miller, O. Fong, K. E. Hirokawa, K. Lathia, C. Rimorin, M. Tieu, R. Larsen, T. Casper, E. Barkan, M. Kroll, S. Parry, N. V. Shapovalova, D. Hirschstein, J. Pendergraft, H. A. Sullivan, T. K. Kim, A. Szafer, N. Dee, P. Groblewski, I. Wickersham, A. Cetin, J. A. Harris, B. P. Levi, S. M. Sunkin, L. Madisen, T. L. Daigle, L. Looger, A. Bernard, J. Phillips, E. Lein, M. Hawrylycz, K. Svoboda, A. R. Jones, C. Koch, H. Zeng, Shared and distinct transcriptomic cell types across neocortical areas, Nature 563, 72–78 (2018).

3. H. Zeng, E. H. Shen, J. G. Hohmann, S. W. Oh, A. Bernard, J. J. Royall, K. J. Glattfelder, S. M. Sunkin, J. A. Morris, A. L. Guillozet-Bongaarts, K. A. Smith, A. J. Ebbert, B. Swanson, L. Kuan, D. T. Page, C. C. Overly, E. S. Lein, M. J. Hawrylycz, P. R. Hof, T. M. Hyde, J. E. Kleinman, A. R. Jones, Large-scale cellular-resolution gene profiling in human neocortex reveals species-specific molecular signatures, Cell 149, 483–496 (2012).

4. N. W. Gouwens, S. A. Sorensen, J. Berg, C. Lee, T. Jarsky, J. Ting, S. M. Sunkin, D. Feng, C. A. Anastassiou, E. Barkan, K. Bickley, N. Blesie, T. Braun, K. Brouner, A. Budzillo, S. Caldejon, T. Casper, D. Castelli, P. Chong, K. Crichton, C. Cuhaciyan, T. L. Daigle, R. Dalley, N. Dee, T. Desta, S.-L. Ding, S. Dingman, A. Doperalski, N. Dotson, T. Egdorf, M. Fisher, R. A. de Frates, E. Garren, M. Garwood, A. Gary, N. Gaudreault, K. Godfrey, M. Gorham, H. Gu, C. Habel, K. Hadley, J. Harrington, J. A. Harris, A. Henry, D. Hill, S. Josephsen, S. Kebede, L. Kim, M. Kroll, B. Lee, T. Lemon, K. E. Link, X. Liu, B. Long, R. Mann, M. McGraw, S. Mihalas, A. Mukora, G. J. Murphy, L. Ng, K. Ngo, T. N. Nguyen, P. R. Nicovich, A. Oldre, D. Park, S. Parry, J. Perkins, L. Potekhina, D. Reid, M. Robertson, D. Sandman, M. Schroedter, C. Slaughterbeck, G. Soler-Llavina, J. Sulc, A. Szafer, B. Tasic, N. Taskin, C. Teeter, N. Thatra, H. Tung, W. Wakeman, G. Williams, R. Young, Z. Zhou, C. Farrell, H. Peng, M. J. Hawrylycz, E. Lein, L. Ng, A. Arkhipov, A. Bernard, J. W. Phillips, H. Zeng, C. Koch, Classification of electrophysiological and morphological neuron types in the mouse visual cortex, Nat. Neurosci. 22, 1182–1195 (2019).

5. R. D. Hodge, T. E. Bakken, J. A. Miller, K. A. Smith, E. R. Barkan, L. T. Graybuck, J. L. Close, B. Long, N. Johansen, O. Penn, Z. Yao, J. Eggermont, T. Höllt, B. P. Levi, S. I. Shehata, B. Aevermann, A. Beller, D. Bertagnolli, K. Brouner, T. Casper, C. Cobbs, R. Dalley, N. Dee, S.-L. Ding, R. G. Ellenbogen, O. Fong, E. Garren, J. Goldy, R. P. Gwinn, D. Hirschstein, C. D. Keene, M. Keshk, A. L. Ko, K. Lathia, A. Mahfouz, Z. Maltzer, M. McGraw, T. N. Nguyen, J. Nyhus, J. G. Ojemann, A. Oldre, S. Parry, S. Reynolds, C. Rimorin, N. V. Shapovalova, S. Somasundaram, A. Szafer, E. R. Thomsen, M. Tieu, G. Quon, R. H. Scheuermann, R. Yuste, S. M. Sunkin, B. Lelieveldt, D. Feng, L. Ng, A. Bernard, M. Hawrylycz, J. W. Phillips, B. Tasic, H. Zeng, A. R. Jones, C. Koch, E. S. Lein, Conserved cell types with divergent features in human versus mouse cortex, Nature 573, 61–68 (2019).

6. H.-J. Suk, E. S. Boyden, I. van Welie, Advances in the automation of whole-cell patch clamp technology, J. Neurosci. Methods 326, 108357 (2019).

7. Y. Peng, F. X. Mittermaier, H. Planert, U. C. Schneider, H. Alle, J. R. P. Geiger, High-throughput microcircuit analysis of individual human brains through next-generation multineuron patch-clamp, doi:10.1101/639328.

8. S. B. Kodandaramaiah, G. L. Holst, I. R. Wickersham, A. C. Singer, G. T. Franzesi, M. L. McKinnon, C. R. Forest, E. S. Boyden, Assembly and operation of the autopatcher for automated intracellular neural recording in vivo, Nat. Protoc. 11, 634–654 (2016).

9. S. B. Kodandaramaiah, G. T. Franzesi, B. Y. Chow, E. S. Boyden, C. R. Forest, Automated whole-cell patch-clamp electrophysiology of neurons in vivo, Nat. Methods 9, 585–587 (2012).

10. S. B. Kodandaramaiah, thesis, Georgia Institute of Technology (2012).

11. H.-J. Suk, I. van Welie, S. B. Kodandaramaiah, B. Allen, C. R. Forest, E. S. Boyden, Closed-Loop Real-Time Imaging Enables Fully Automated Cell-Targeted Patch-Clamp Neural Recording In Vivo, Neuron 96, 244–245 (2017).

12. B. Long, L. Li, U. Knoblich, H. Zeng, H. Peng, 3D Image-Guided Automatic Pipette Positioning for Single Cell Experiments in vivo, Sci. Rep. 5, 18426 (2015).

13. L. A. Annecchino, A. R. Morris, C. S. Copeland, O. E. Agabi, P. Chadderton, S. R. Schultz, Robotic Automation of In Vivo Two-Photon Targeted Whole-Cell Patch-Clamp Electrophysiology, Neuron 95, 1048–1055.e3 (2017).

14. N. S. Desai, J. J. Siegel, W. Taylor, R. A. Chitwood, D. Johnston, MATLAB-based automated patch-clamp system for awake behaving mice, J. Neurophysiol. 114, 1331–1345 (2015).

15. W. A. Stoy, I. Kolb, G. L. Holst, Y. Liew, A. Pala, B. Yang, E. S. Boyden, G. B. Stanley, C. R. Forest, Robotic navigation to subcortical neural tissue for intracellular electrophysiology in vivo Journal of Neurophysiology 118, 1141–1150 (2017).

16. I. Kolb, W. A. Stoy, E. B. Rousseau, O. A. Moody, A. Jenkins, C. R. Forest, Cleaning patch-clamp pipettes for immediate reuse, Sci. Rep. 6, 35001 (2016).

17. I. Kolb, C. R. Landry, M. C. Yip, C. F. Lewallen, W. A. Stoy, J. Lee, A. Felouzis, B. Yang, E. S. Boyden, C. J. Rozell, C. R. Forest, PatcherBot: a single-cell electrophysiology robot for adherent cells and brain slices, J. Neural Eng. 16, 046003 (2019).

18. R. Perin, H. Markram, A computer-assisted multi-electrode patch-clamp system, J. Vis. Exp., e50630 (2013).

19. S. B. Kodandaramaiah, F. J. Flores, G. L. Holst, A. C. Singer, X. Han, E. N. Brown, E. S. Boyden, C. R. Forest, Multi-neuron intracellular recording in vivo via interacting autopatching robots, Elife 7 (2018), doi:10.7554/eLife.24656.

20. L. Li, B. Ouellette, W. A. Stoy, E. J. Garren, T. L. Daigle, C. R. Forest, C. Koch, H. Zeng, A robot for high yield electrophysiology and morphology of single neurons in vivo, Nat. Commun. 8, 15604 (2017).

21. K. Koos, J. Molnár, P. Horvath, Pipette Hunter: Patch-Clamp Pipette Detection Image Analysis, 172–183 (2017).

22. Q. Wu 吴秋雫, I. Kolb, B. M. Callahan, Z. Su, W. Stoy, S. B. Kodandaramaiah, R. Neve, H. Zeng, E. S. Boyden, C. R. Forest, A. A. Chubykin, Integration of autopatching with automated pipette and cell detection in vitro, J. Neurophysiol. 116, 1564–1578 (2016).

23. E. Moen, D. Bannon, T. Kudo, W. Graf, M. Covert, D. Van Valen, Deep learning for cellular image analysis Nature Methods 16, 1233–1246 (2019).

24. I. Suleymanova, T. Balassa, S. Tripathi, C. Molnar, M. Saarma, Y. Sidorova, P. Horvath, A deep convolutional neural network approach for astrocyte detection Scientific Reports 8 (2018), doi:10.1038/s41598-018-31284-x.

25. Allen Institute for Brain Science, Allen Cell Types Database Allen Brain Atlas (available at http://help.brain-map.org/display/celltypes).

26. A. E. Carpenter, T. R. Jones, M. R. Lamprecht, C. Clarke, I. H. Kang, O. Friman, D. A. Guertin, J. H. Chang, R. A. Lindquist, J. Moffat, P. Golland, D. M. Sabatini, CellProfiler: image analysis software for identifying and quantifying cell phenotypes, Genome Biol. 7, R100 (2006).

27. C. Sommer, C. Straehle, U. Kothe, F. A. Hamprecht, Ilastik: Interactive learning and segmentation toolkit 2011 IEEE International Symposium on Biomedical Imaging: From Nano to Macro (2011), doi:10.1109/isbi.2011.5872394.

28. C. Tomasi, T. Kanade, Detection and Tracking of Point Features (1991).

29. J. Shi, Tomasi, Good features to track Proceedings of IEEE Conference on Computer Vision and Pattern Recognition CVPR-94 (1994), doi:10.1109/cvpr.1994.323794.

30. K. Koos, J. Molnár, L. Kelemen, G. Tamás, P. Horvath, DIC image reconstruction using an energy minimization framework to visualize optical path length distribution, Sci. Rep. 6, 30420 (2016).

31. R. Yang, K. W. C. Lai, N. Xi, J. Yang, Development of automated patch clamp system for electrophysiology 2013 IEEE International Conference on Robotics and Biomimetics (ROBIO) (2013), doi:10.1109/robio.2013.6739793.

32. H. Markram, E. Muller, S. Ramaswamy, M. W. Reimann, M. Abdellah, C. A. Sanchez, A. Ailamaki, L. Alonso-Nanclares, N. Antille, S. Arsever, G. A. A. Kahou, T. K. Berger, A. Bilgili, N. Buncic, A. Chalimourda, G. Chindemi, J.-D. Courcol, F. Delalondre, V. Delattre, S. Druckmann, R. Dumusc, J. Dynes, S. Eilemann, E. Gal, M. E. Gevaert, J.-P. Ghobril, A. Gidon, J. W. Graham, A. Gupta, V. Haenel, E. Hay, T. Heinis, J. B. Hernando, M. Hines, L. Kanari, D. Keller, J. Kenyon, G. Khazen, Y. Kim, J. G. King, Z. Kisvarday, P. Kumbhar, S. Lasserre, J.-V. Le Bé, B. R. C. Magalhães, A. Merchán-Pérez, J. Meystre, B. R. Morrice, J. Muller, A. Muñoz-Céspedes, S. Muralidhar, K. Muthurasa, D. Nachbaur, T. H. Newton, M. Nolte, A. Ovcharenko, J. Palacios, L. Pastor, R. Perin, R. Ranjan, I. Riachi, J.-R. Rodríguez, J. L. Riquelme, C. Rössert, K. Sfyrakis, Y. Shi, J. C. Shillcock, G. Silberberg, R. Silva, F. Tauheed, M. Telefont, M. Toledo-Rodriguez, T. Tränkler, W. Van Geit, J. V. Díaz, R. Walker, Y. Wang, S. M. Zaninetta, J. DeFelipe, S. L. Hill, I. Segev, F. Schürmann, Reconstruction and Simulation of Neocortical Microcircuitry, Cell 163, 456–492 (2015).

33. Petilla Interneuron Nomenclature Group, G. A. Ascoli, L. Alonso-Nanclares, S. A. Anderson, G. Barrionuevo, R. Benavides-Piccione, A. Burkhalter, G. Buzsáki, B. Cauli, J. Defelipe, A. Fairén, D. Feldmeyer, G. Fishell, Y. Fregnac, T. F. Freund, D. Gardner, E. P. Gardner, J. H. Goldberg, M. Helmstaedter, S. Hestrin, F. Karube, Z. F. Kisvárday, B. Lambolez, D. A. Lewis, O. Marin, H. Markram, A. Muñoz, A. Packer, C. C. H. Petersen, K. S. Rockland, J. Rossier, B. Rudy, P. Somogyi, J. F. Staiger, G. Tamas, A. M. Thomson, M. Toledo-Rodriguez, Y. Wang, D. C. West, R. Yuste, Petilla terminology: nomenclature of features of GABAergic interneurons of the cerebral cortex, Nat. Rev. Neurosci. 9, 557–568 (2008).

34. N. Faragó, á. K. Kocsis, S. Lovas, G. Molnár, E. Boldog, M. Rózsa, V. Szemenyei, E. Vámos, L. I. Nagy, G. Tamás, L. G. Puskás, Digital PCR to determine the number of transcripts from single neurons after patch-clamp recording, Biotechniques 54, 327–336 (2013).

35. N. Faragó, á. K. Kocsis, C. Braskó, S. Lovas, M. Rózsa, J. Baka, B. Kovács, K. Mikite, V. Szemenyei, G. Molnár, A. Ozsvár, G. Oláh, I. Piszár, á. Zvara, A. Patócs, P. Barzó, L. G. Puskás, G. Tamás, Human neuronal changes in brain edema and increased intracranial pressure, Acta Neuropathol Commun 4, 78 (2016).

36. G. Molnár, N. Faragó, á. K. Kocsis, M. Rózsa, S. Lovas, E. Boldog, R. Báldi, É. Csajbók, J. Gardi, L. G. Puskás, G. Tamás, GABAergic neurogliaform cells represent local sources of insulin in the cerebral cortex, J. Neurosci. 34, 1133–1137 (2014).

