## Supplementary Information: Cell Detection System for "Automatic deep learning driven label-free image guided patch clamp system for human and rodent in vitro slice physiology"

*Supplementary Information*

#### Algorithm

The aim of the cell detection system is to automatically detect cells in tissue image stacks and thereafter suggest them for automatic patch clamp recording. We created and labeled image stacks and trained a deep convolutional neural network. The detection is performed on slices of 2D images and the results are extended to the 3D where the refined position of the cells is determined in increased quality. The extension was done by merging overlapping detection bounding boxes over adjacent slices by using their intersection regions. The required overlapping ratio can be set as a parameter. The merging can still happen even if a cell is not detected on every consecutive slice. If one slice does not contain a detection in the position of a cell but the previous and the subsequent do then the slice in question is assumed to contain the object and it will also be included. The merging algorithm compensates for the noise sensitivity of the detection.

#### Labeling

Our software provides a labeling tool (see Supplementary Information: Software Usage) that offers a platform to generate an applicable annotated dataset. Field experts labeled 6344 cells on 265 stacks which took about 132 hours. The annotation procedure consisted of putting bounding boxes around the recognized cells over multiple slices in the stack. The acquired labeled training set was converted into the format of the deep learning framework we used.

#### Training process

The number of deep learning frameworks had increased in the past few years. We chose the *Caffe* [1] deep learning framework because of its running time and support of both python and Matlab programming languages. The framework allows adding, removal and modification of the elements of deep learning architectures which was a requirement.

As an extension of Caffe, the NVIDIA's Deep Learning GPU Training System (DIGITS [2]) allows the non-advanced deep learning users to evaluate the finest training at ease. DIGITS also includes a pre-defined neural network architecture called DetectNet [3]. DetectNet can be split into training and validation sections and these can be further divided into three and two subsections, respectively. The training part contains the data augmentation, the fully connected network (GoogLeNet [4]) and the loss functions, while the bounding box clustering and the mean average precision (mAP) statistics are the elements of the validation part.

DIGITS was used to train DetectNet. The solver used for the training process was Adaptive Moment Estimation (ADAM) [5]. The learning rate was 1e-5 with a fixed learning rate policy. The pre-trained weights of the ImageNet dataset were used for the initialization of GoogLeNet to speed up the training process. The training was performed on a PC with Intel Core i7-4770K 3.5 GHz CPU, 16 GB memory and an NVIDIA Titan Xp graphics card. The number of epochs was 2500 which took 6 days and 15 hours.

#### Evaluation

To evaluate the performance of the proposed framework we measured precision, recall and F1 score of two different annotators. The object matching between the ground truth data and the detection results was done manually. This resulted in better matching when multiple ground truth bounding boxes were close to each other or even overlapped. A detection was considered to be correct (true positive, TP) when it significantly overlapped with a ground truth bounding box at least on one image slice in the stack. Otherwise, it was treated as a false positive (FP) detection. Ground truth objects not paired with a detection were treated as false negatives (FN). Based on these aspects the detection accuracy was calculated as precision P=TP/(TP+FP), recall R=TP/(TP+FN), and F1 score=2*P*R/(P+R).
For the final result, we evaluated four different comparisons. First, we determined intra- and inter-expert accuracies. The cell-detection task proved to be a complex issue as the two annotators reached 0.771 (precision = 83.12%, recall = 71.91%) and 0.778 F1-score (precision = 87.5%, recall = 70%), respectively. To compare the experts, the inter-expert accuracy was measured which resulted in 0.752 F1-score (precision = 81.54%, recall = 69.74%). For computing all the accuracies, we used the first annotation set of the first annotator as a ground truth. As the final test, we compared the ground truth to the bounding boxes predicted by our method. This outperformed both of the annotators with 0.835 F1-score (precision = 81.72%, recall = 85.39%) (Fig. 1).

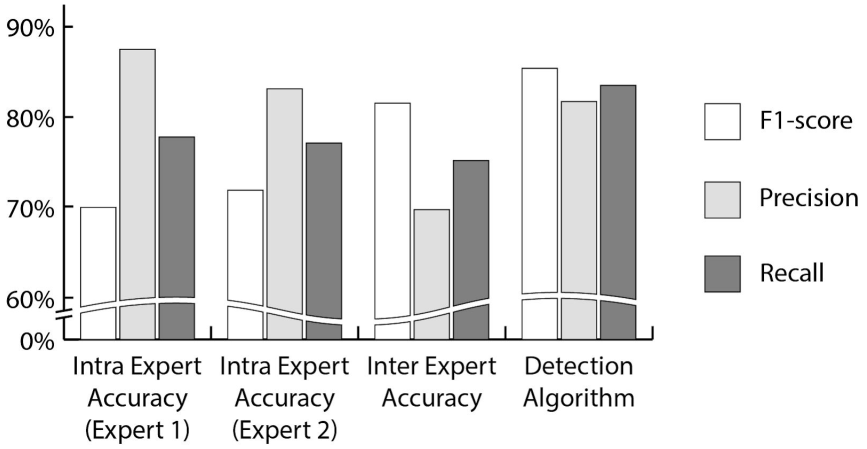


Figure 1: **Evaluation results.**

### Prediction on a server

We developed a tool to perform cell detection on a server computer which tool we refer to as RemoteWorker. The microscope computers operate as clients which send the image to the server and download the detection result, thus they share a GPU. The tool can easily be configured to run any Matlab script on a different computer. The code used for remote cell detection is included in the software and can be selected in the configuration file. Below an example shows how to set up both the client and server code to add 5 to a number.
The server-side script should register the supported commands as follows:

```
s = RemoteWorker.Server();
s.registerCommand('addFive', @(header,data) data+5);
s.run();
```

The client sends a job, polls if the results are ready and then downloads the return value:

```
c = RemoteWorker.Worker();
c.host = '127.0.0.1';
c.sendJob('addFive', 1);
```

```
while ~c.isJobReady()
    pause(0.01);
end
result = c.downloadResult(); % returns 6
```

### References

[1] Jia, Y. et al. Caffe: Convolutional Architecture for Fast Feature Embedding. Proceedings of the 22nd ACM international conference on Multimedia , 675–678, (2014)
[2] Yeager, Luke, et al. DIGITS: the deep learning GPU training system. ICML 2015 AutoML Workshop, (2015)
[3] Tao, Andrew, Jon Barker, and Sriya Sarathy. "Detectnet: Deep neural network for object detection in digits." Parallel Forall 4 (2016).
[4] Szegedy, C., Liu, W., Jia, Y., et al. Going Deeper with Convolutions. Proceedings of the IEEE conference on computer vision and pattern recognition , 1-9, (2015)
[5] Kingma, D. P., & Ba, J. L. Adam: a Method for Stochastic Optimization. International Conference on Learning Representations , 1–13, (2015)
