## Supplementary Information: Cell Tracking System for "Automatic deep learning driven label-free image guided patch clamp system for human and rodent in vitro slice physiology"

*Supplementary Information*

#### Introduction

The target cell during the patch clamping process can shift as the pipette is being pushed towards it. The shift can occur in any direction in the tissue and based on our experiments it is usually between 3 and 10 µm (6.98 ± 3.91 µm, n=19). Tracking the cell under the microscope is a challenging task because a 3-dimensional tracking is required with a 2-dimensional label-free modality. The developed online tracking system has two parts. One part performs lateral tracking in the XY plane while the other part tracks the cell in the Z dimension. Both parts require a template image of the target cell which is acquired before starting the patch clamp process and when the cell is in focus in the image. The lateral tracking is always performed in the image of the latest focus level. The Z tracking algorithm operates on a small image stack. Acquiring an image stack is time-consuming as the objective has to be moved physically to different focus levels. Therefore, when Z tracking is being performed, the pipette movement is paused to ensure that the cell is not pushed meanwhile.

#### Lateral cell tracking

The lateral tracker is the Kanade-Lucas-Tomasi (KLT) feature tracker algorithm. Feature points are detected by the minimum eigenvalue algorithm in a window in the template image [1,2]. The algorithm is computationally inexpensive and it is performed often (usually per second) except when Z tracking is in progress. Supplementary Video 2 shows the performance of the lateral tracker. The white '+' markers at the beginning of the examples in the video show the detected feature points on which the tracker is based. Our experimental verification shows that in 95% of the cases the algorithm was able to track single cells for more than 100 frames, even in case of extreme motion (>3 µm/frame).
The default window size for the tracker was 241x241 pixels which is enough to cover almost every cell (except rarely large human cells). The feature points are detected using a 5x5 filter. If the number of valid feature points significantly drops, for example, due to a sudden movement of the cell in the Z direction, the points are reinitialized. A significant drop of feature points is detected when the number of valid points is less than 10% of their original number or less than or equal to 10.

#### Tracking in Z dimension

Tracking in the Z dimension is based on focus detection algorithms. As a major principle, we assume that an object which is in focus has sharp edges, that cause large differences in pixel intensities. In the beginning, a small image stack is acquired around the last known position of the target cell. The template image is compared to every slice of the stack by calculating the absolute value of the difference of the standard deviations of the images in a small window. The position of the window is determined by the lateral tracker. The maximums of the computed values are selected for the images below and above the middle element of the stack. The middle slice corresponds to the last known focus position. Then the minimum of these three values, the two maximums and the one of the middle slice, determine the direction of the shift of the cell. Fig. 1 shows examples of the computed values and the related images based on which the decisions are made. If the lowest value is of the middle slice, the cell has not moved. If the cell has moved below its previous position then the minimum value will be that of the elements below the middle slice. Similarly, we can determine if the cell has moved up. Although this approach does not specify the displacement value only the direction of it, the tracking is reliable because the cells do not move rapidly.

The stack size used for the tracking, based on empirical tests, consists of 7 slices by default, thus the middle element is the 4th. The data shown in Fig. 1 are calculated from bigger image stacks, consisting of 60-100 slices. The cells shown were selected from the results of the cell detection system. This allowed showing more values than just 7 for demonstration in the figure.

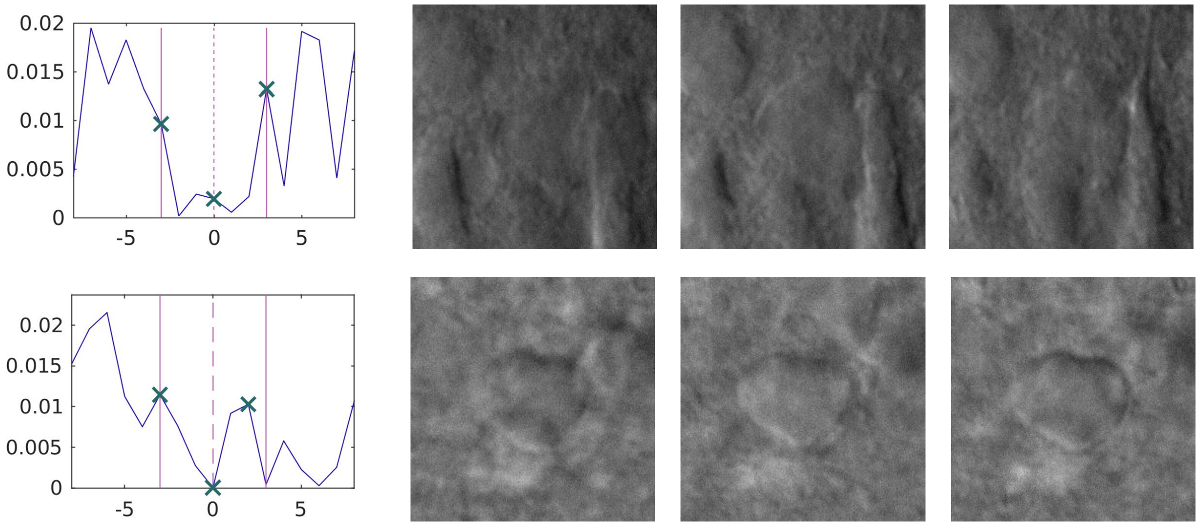


Figure 1: **Examples of the Z tracking system.** The plots in the first column show the calculated values used for determining the direction of the shift. The X-axis shows the index difference from the middle slice. The corresponding image regions of the markers in the plots are shown in the next three columns. The final decision whether the cell has shifted is based on these values and images. The lines at ±3 in the X-axis indicate the region which is used in the calculations, while the values outside this interval are for demonstration purposes.

### References

[1] Bruce D. Lucas and Takeo Kanade. An Iterative Image Registration Technique with an Application to Stereo Vision. International Joint Conference on Artificial Intelligence, pages 674–679, 1981.

[2] Carlo Tomasi and Takeo Kanade. Detection and Tracking of Point Features. Carnegie Mellon University Technical Report CMU-CS-91-132, April 1991.
