## Supplementary Information: Pipette Detection System for "Automatic deep learning driven label-free image guided patch clamp system for human and rodent in vitro slice physiology"

*Supplementary Information*

#### 1 Introduction

The pipette detection pipeline consists of three steps. First, when the pipette is visible in the image, an image stack is acquired. Ideally, the pipette should cover most of the image and should be nearly in focus. Moreover, the tip should be near the sample so that light conditions during detection are similar to those during the movement of the pipette in the sample. Then a fast initialization algorithm is run on the acquired stack. The initialization is based on a moving line profile in the image and is detailed in Section 2. Finally, the result of the initialization is refined using the main tip detection algorithm referred to as Pipette Hunter 3D (PH3D). The algorithm is the 3-dimensional extension of the base Pipette Hunter model [1] and the derivations are detailed in Section 3. The energy function is minimized using the variational method and gradient descent. Section 4 describes the performed quality measurement and comparison of the algorithms.

#### 2 Line profile estimation

An initialization is advantageous to the main detection algorithm as variational frameworks are known to be robust but require many iterations and the result is sensitive to the starting point. The developed initialization algorithm operates on minimum intensity projected (MIP) images of the stack containing the pipette. Two independent runs are required on different projection images (usually Z and X projections) to get a 3D point estimation.

The initialization algorithm estimates the tip position by analyzing a line profile in the MIP image. First, it is assumed that the orientation of the pipette is known either by setting this value manually or using the value set by the pipette calibration step. Then a line is placed in front of the pipette on the edge of the image and rotated such that it is perpendicular to the orientation of the pipette. Then the line profile is analyzed in those positions where it covers the image. Since the image is often noisy we compute the 90th percentile of the pixel values from the line profile which will be a reference value. If the profile includes at least one value that is relevantly smaller than the reference value it is considered to be caused by the edge of the pipette. The relevantly smaller value is defined as a 40% intensity drop but it can be set as a parameter to the algorithm. If more than one relevant values are present the smallest is chosen. Subsequent to this the position in the image of the chosen relevant small value is computed from the parameters of the line, the algorithm terminates and outputs this value as the position of the pipette tip. If no such value is present in the profile then the line is pushed towards the pipette tip by one pixel distance and the algorithm repeats from analyzing the line's profile. The different steps of the algorithm are presented in Fig. 1. The quality measurement of this initialization step is described in Section 4.

#### 3 Pipette Hunter 3D

The **notations** used throughout this section are listed below:

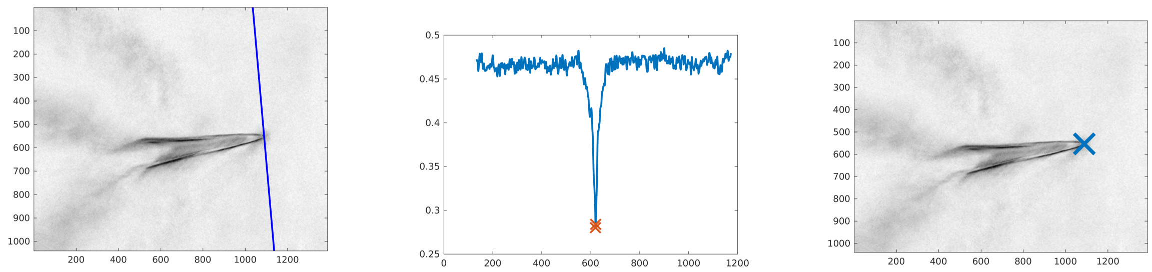


a b c

Figure 1: Initialization using line profile estimation. **a:** A line aligned perpendicularly to the orientation of the pipette is pushed against the pipette from the edge of the image. **b:** The line profile of the image where the first relevant intensity drop was found. **c:** The position of the intensity drop in the line profile is converted back to a position in the image.

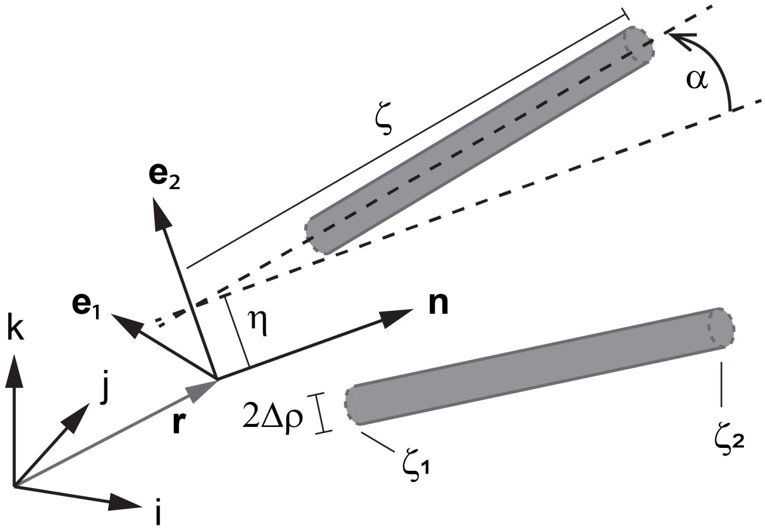


Figure 2: The 8 DOF pipette detection mechanism.

- tensors (including vectors) are distinguished from scalars or coordinates using bold letters
- we use different notations for tensors and their representations: representation $[\mathbf{A}]$ or coordinate matrix (w.r.t. some coordinate system) encased in brackets is associated with tensor $\mathbf{A}$
- for dot (scalar) and cross products between vectors, we use operators "$\cdot$" and "$\times$" respectively
- dot operation is used in a broader "contraction" sense: (single-) dot product reduces the order of the respective tensor product by two; examples a) let $\mathbf{v}$, $\mathbf{w}$ be arbitrary vectors (tensors of 1st order), their tensor (dyadic) product is a second order tensor, then $\mathbf{v} \cdot \mathbf{w}$ is scalar (tensor of 0-th order) b) $\mathbf{w} = \mathbf{A} \cdot \mathbf{v}$ where $\mathbf{A}$ is second order tensor and $\mathbf{v}$ is vector, then $\mathbf{w}$ is also a vector c) $\mathbf{C} = \mathbf{A} \cdot \mathbf{B}$ all tensors of order two; between the representations, these operations are performed using the usual matrix-vector and matrix-matrix multiplications (e.g. $[\mathbf{w}] = [\mathbf{A}]\,[\mathbf{v}]$, $[\mathbf{C}] = [\mathbf{A}]\,[\mathbf{B}]$)

- to denote the gradient of a scalar field $I(\mathbf{r})$ (where $\mathbf{r}$ is a position vector w.r.t. some coordinate system) we use either $\frac{\partial I}{\partial \mathbf{r}}$ or the "right gradient" notation $I\nabla$.

Here, we define the energy and derive its gradient for the two-wand mechanism designed to detect the pipette (see Fig.2). The assumptions are as follows:

- original configuration: the moving local frame $(\mathbf{e}_1, \mathbf{e}_2, \mathbf{n})$ with coordinates $\xi^1, \xi^2, \xi^3$ is aligned with the standard basis $(\mathbf{i}, \mathbf{j}, \mathbf{k})$ with coordinates denoted by $x^1, x^2, x^3$
- the axes of the wands are situated on the plane spanned by $\mathbf{e}_2, \mathbf{n}$, satisfying the equation $\xi^2 = \pm\eta^1 \pm \xi^3 \tan\alpha^1$
- the symmetrical arrangement of the wands is retained during the iterative approaching of the energy minimum
- $\zeta$ is the distance along the wand axes measured from the plane spanned by $\mathbf{e}_1, \mathbf{e}_2$, $\rho$ is the distance to the wand axes and $\phi$ is the angle around it, measured from $\mathbf{e}_1$ in the plane $\mathbf{e}_1, \mathbf{e}_2$.

The wand positions are completely determined by (together with the symmetry condition above)

- the position vector $\mathbf{r}(x^1, x^2, x^3) = x^1\mathbf{i} + x^2\mathbf{j} + x^3\mathbf{k}$ that designates the origin (or pivot point) of the local frame $(\mathbf{e}_1, \mathbf{e}_2, \mathbf{n})$
- the rotation of the local frame about its pivot point (we parameterize the space of rotations with proper Euler angles $\varphi^1, \varphi^2, \varphi^3$, using extrinsic $z - y - z$ scheme)
- the coordinates $\eta^1, \alpha^1$ which are given relative to the moving local frame
- the (fixed parameter) values $\zeta_1, \zeta_2$ representing the start and the endpoints of the wands along their axes.

The system has therefore 8 degrees of freedom (DOF) given by the variables having upper indices in the list above: $x^1, x^2, x^3, \varphi^1, \varphi^2, \varphi^3, \eta^1, \alpha^1$.

The local coordinates of a point w.r.t. the local frame $(\mathbf{e}_1, \mathbf{e}_2, \mathbf{n})$ inside the cylinder-shaped wands, using cylindrical coordinates $\rho, \phi, \zeta$ are given by:

$$\begin{aligned} \xi^1(\rho,\phi,\zeta) &= \rho\cos\phi \\ \xi^2(\rho,\phi,\zeta) &= \pm\left(\eta^1 + \zeta\sin\alpha^1\right) + (\rho\sin\phi)\cos\alpha^1 \\ \xi^3(\rho,\phi,\zeta) &= \zeta\cos\alpha^1 \mp (\rho\sin\phi)\sin\alpha^1, \end{aligned} \tag{1}$$

$\rho \in [0, \triangle\rho]$, $\zeta \in [\zeta_1, \zeta_2]$, $\phi \in [0, 2\pi)$, then the internal point of wands $\mathbf{P}\left(x^i, \varphi^k, \eta^1, \alpha^1\right) = \mathbf{r} + \xi^1\mathbf{e}_1 + \xi^2\mathbf{e}_2 + \xi^3\mathbf{n}$, where $i$, $k = 1, 2, 3$ is identified by the formula:

$$\begin{aligned} \mathbf{P}_\pm = \mathbf{r}\left(x^i\right) & \\ &\pm \eta^1\mathbf{e}_2\left(\varphi^k\right) + \zeta\left(\cos\alpha^1\mathbf{n}\left(\varphi^k\right) \pm \sin\alpha^1\mathbf{e}_2\left(\varphi^k\right)\right) \\ &+ \rho\left[\sin\phi\left(\cos\alpha^1\mathbf{e}_2\left(\varphi^k\right) \mp \sin\alpha^1\mathbf{n}\left(\varphi^k\right)\right) + \cos\phi\mathbf{e}_1\left(\varphi^k\right)\right]. \end{aligned} \tag{2}$$

Note that in the expression (2), the second line identifies the points of the wand axes. Now we are ready to define the energy of the system as the integral of the voxel-intensity over the volume occupied with the wands:

$$E\left(x^i,\varphi^k,\eta^1,\alpha^1\right) \doteq \int_V \left[I\left(\mathbf{P}_+\right)+I\left(\mathbf{P}_+\right)\right]dV$$
$$= \int_{\zeta_1}^{\zeta_2}\int_0^{2\pi}\int_0^{\triangle\rho}\left[I\left(\mathbf{P}_+\right)+I\left(\mathbf{P}_+\right)\right]\rho d\rho d\phi d\zeta. \quad (3)$$

The local coordinates as the functions of the variables of the integration $\rho,\phi,\zeta$ are given by (1). Energy (3) has its minimal value wherever its derivatives w.r.t. the variables defining the system DOF are all zero. The list of the required derivatives is:

$$\begin{aligned}
\frac{\partial E}{\partial \mathbf{r}} &= \int_V \left[I\nabla\left(\mathbf{P}_+\right)+I\nabla\left(\mathbf{P}_-\right)\right]dV \\
\frac{\partial E}{\partial \varphi^k} &= \int_V \left[I\nabla\left(\mathbf{P}_+\right)\cdot\frac{\partial \mathbf{P}_+}{\partial\varphi^k}+I\nabla\left(\mathbf{P}_-\right)\cdot\frac{\partial \mathbf{P}_-}{\partial\varphi^k}\right]dV \\
\frac{\partial E}{\partial \eta^1} &= \int_V \left[I\nabla\left(\mathbf{P}_+\right)\cdot\frac{\partial \mathbf{P}_+}{\partial\eta^1}+I\nabla\left(\mathbf{P}_-\right)\cdot\frac{\partial \mathbf{P}_-}{\partial\eta^1}\right]dV \qquad (4)\\
\frac{\partial E}{\partial \alpha^1} &= \int_V \left[I\nabla\left(\mathbf{P}_+\right)\cdot\frac{\partial \mathbf{P}_+}{\partial\alpha^1}+I\nabla\left(\mathbf{P}_-\right)\cdot\frac{\partial \mathbf{P}_-}{\partial\alpha^1}\right]dV\,.
\end{aligned}$$

Below we examine the integrands of (4) line by line.

**1st line:** The components of $\frac{\partial E}{\partial \mathbf{r}}$ are the derivatives w.r.t. the coordinates of the origin of the local frame in the world coordinate system: $\frac{\partial E}{\partial x^i}=\frac{\partial E}{\partial \mathbf{r}}\cdot\mathbf{l}$, $\mathbf{l}\in\{\mathbf{i},\mathbf{j},\mathbf{k}\}$. $I\nabla$ is the 'right' gradient (represented by row vector) of the image function $I\left(x^1,x^2,x^3\right)$, "$\cdot$" stands for the dot (scalar) product of vectors.

**2nd line:** Since the local coordinates (1) are known for all values of integration variables $\rho\in[0,\triangle\rho]$, $\zeta\in[\zeta_1,\zeta_2]$, $\phi\in[0,2\pi)$, we wish to express the quantities that occur in the list of derivatives (4) with these local coordinates. According to (2), only the derivatives of the rotated local frame basis vectors w.r.t. the Euler angles are required. The columns of the representing matrix of the rotation $\mathbf{R}=\mathbf{R}_{\varphi^1}\cdot\mathbf{R}_{\varphi^2}\cdot\mathbf{R}_{\varphi^3}$ is calculated as the multiplication of the matrices of the elementary rotations (applied from the left to column vectors)

$$[\mathbf{R}]=\begin{bmatrix} c_1c_2c_3-s_1s_3 & -c_3s_1-c_1c_2s_3 & c_1s_2 \\ c_1s_3+c_2c_3s_1 & c_1c_3-c_2s_1s_3 & s_1s_2 \\ -c_3s_2 & s_2s_3 & c_2 \end{bmatrix}=\begin{bmatrix}\mathbf{e}_1 & \mathbf{e}_2 & \mathbf{n}\end{bmatrix}, \qquad (5)$$

its columns contain the coordinates of the basis vectors of the local system $(\mathbf{e}_1,\mathbf{e}_2,\mathbf{n})$ expressed in the standard basis (the first two angles $\varphi^1$ and $\varphi^2$ designate the azimuthal and polar coordinates of $\mathbf{n}$ respectively whilst $\varphi^3$ represents the spin of the plane spanned by $\mathbf{e}_1,\mathbf{e}_2$). Here we use the usual abbreviations $s_i\equiv\sin\varphi^i$, $c_k\equiv\cos\varphi^k$, $i$, $k=1,2,3$ for concise writing. The derivatives are determined by direct calculation, the results can be expressed by the angles and the local basis vectors as:

$$
\begin{array}{lll}
\dfrac{\partial \mathbf{e}_1}{\partial \varphi^1} = c_2\mathbf{e}_2 - s_2 s_3 \mathbf{n} & \dfrac{\partial \mathbf{e}_1}{\partial \varphi^2} = -c_3\mathbf{n} & \dfrac{\partial \mathbf{e}_1}{\partial \varphi^3} = \mathbf{e}_2 \\[2ex]
\dfrac{\partial \mathbf{e}_2}{\partial \varphi^1} = -c_2\mathbf{e}_1 - s_2 c_3 \mathbf{n} & \dfrac{\partial \mathbf{e}_2}{\partial \varphi^2} = s_3\mathbf{n} & \dfrac{\partial \mathbf{e}_2}{\partial \varphi^3} = -\mathbf{e}_1 \\[2ex]
\dfrac{\partial \mathbf{n}}{\partial \varphi^1} = s_2\left(s_3\mathbf{e}_1 + c_3\mathbf{e}_2\right) & \dfrac{\partial \mathbf{n}}{\partial \varphi^2} = c_3\mathbf{e}_1 - s_3\mathbf{e}_2 & \dfrac{\partial \mathbf{n}}{\partial \varphi^3} = \mathbf{0}.
\end{array}
\tag{6}
$$

Note that these derivatives are of constant length vectors - the unit-length local basis vectors - hence lying in their perpendicular plane (e.g. $\frac{\partial \mathbf{e}_1}{\partial \varphi^k} \in \mathit{span}\left\{\mathbf{e}_2,\, \mathbf{n}\right\}$). From (6), for the point $\mathbf{P} = \mathbf{r} + \xi^1\mathbf{e}_1 + \xi^2\mathbf{e}_2 + \xi^3\mathbf{n}$, $\mathbf{P} \in \left\{\mathbf{P}_+, \mathbf{P}_-\right\}$ the partial derivative expressions: $I\nabla\left(\mathbf{P}\right) \cdot \frac{\partial \mathbf{P}}{\partial \varphi^k}$ (the integrands) w.r.t. the Euler angles $\varphi^1, \varphi^2, \varphi^3$ are calculated such that

$$
\begin{aligned}
I\nabla\left(\mathbf{P}\right) \cdot \frac{\partial \mathbf{P}}{\partial \varphi^1} &= \left(\xi^3 s_2 s_3 - \xi^2 c_2\right)\left(I\nabla \cdot \mathbf{e}_1\right) \\
&\quad + \left(\xi^3 s_2 c_3 + \xi^1 c_2\right)\left(I\nabla \cdot \mathbf{e}_2\right) - \left(\xi^1 s_2 s_3 + \xi^2 s_2 c_3\right)\left(I\nabla \cdot \mathbf{n}\right) \\
I\nabla\left(\mathbf{P}\right) \cdot \frac{\partial \mathbf{P}}{\partial \varphi^2} &= \xi^3 c_3\left(I\nabla \cdot \mathbf{e}_1\right) - \xi^3 s_3\left(I\nabla \cdot \mathbf{e}_2\right) + \left(\xi^2 s_3 - \xi^1 c_3\right)\left(I\nabla \cdot \mathbf{n}\right) \\
I\nabla\left(\mathbf{P}\right) \cdot \frac{\partial \mathbf{P}}{\partial \varphi^3} &= -\xi^2\left(I\nabla \cdot \mathbf{e}_1\right) + \xi^1\left(I\nabla \cdot \mathbf{e}_2\right),
\end{aligned}
\tag{7}
$$

where the image gradient is decomposed in the local system as

$$
I\nabla = \left(I\nabla \cdot \mathbf{e}_1\right)\mathbf{e}_1 + \left(I\nabla \cdot \mathbf{e}_2\right)\mathbf{e}_2 + \left(I\nabla \cdot \mathbf{n}\right)\mathbf{n} \tag{8}
$$

and the wand points are identified with equations (1). To gain deeper insight it is worth introducing the pseudovector $\mathbf{M} \doteq \left(\mathbf{P} - \mathbf{r}\right) \times I\nabla$ - the "torque" - w.r.t. the pivot point of the Euler angles, that is the origin of the local basis $\left(\mathbf{e}_1, \mathbf{e}_2, \mathbf{n}\right)$:

$$
\begin{aligned}
\mathbf{M} &= \left(\xi^1\mathbf{e}_1 + \xi^2\mathbf{e}_2 + \xi^3\mathbf{n}\right) \times \left[\left(I\nabla \cdot \mathbf{e}_1\right)\mathbf{e}_1 + \left(I\nabla \cdot \mathbf{e}_2\right)\mathbf{e}_2 + \left(I\nabla \cdot \mathbf{n}\right)\mathbf{n}\right] \\
&= \left[\xi^2\left(I\nabla \cdot \mathbf{n}\right) - \xi^3\left(I\nabla \cdot \mathbf{e}_2\right)\right]\mathbf{e}_1 \\
&\quad + \left[\xi^3\left(I\nabla \cdot \mathbf{e}_1\right) - \xi^1\left(I\nabla \cdot \mathbf{n}\right)\right]\mathbf{e}_2 \\
&\quad + \left[\xi^1\left(I\nabla \cdot \mathbf{e}_2\right) - \xi^2\left(I\nabla \cdot \mathbf{e}_1\right)\right]\mathbf{n}\,.
\end{aligned}
\tag{9}
$$

With this pseudovector, the constituents of the energy derivatives w.r.t. the Euler angles (7) can be expressed as:

$$
\begin{aligned}
I\nabla\left(\mathbf{P}\right) \cdot \frac{\partial \mathbf{P}}{\partial \varphi^1} &= \mathbf{M} \cdot \left[s_2\left(-c_3\mathbf{e}_1 + s_3\mathbf{e}_2\right) + c_2\mathbf{n}\right] = \mathbf{M} \cdot \mathbf{k} \\
I\nabla\left(\mathbf{P}\right) \cdot \frac{\partial \mathbf{P}}{\partial \varphi^2} &= \mathbf{M} \cdot \left(s_3\mathbf{e}_1 + c_3\mathbf{e}_2\right) = \mathbf{M} \cdot \left(\mathbf{R}_{\varphi^1} \cdot \mathbf{j}\right) \\
I\nabla\left(\mathbf{P}\right) \cdot \frac{\partial \mathbf{P}}{\partial \varphi^3} &= \mathbf{M} \cdot \mathbf{n} = \mathbf{M} \cdot \left(\mathbf{R}_{\varphi^1} \cdot \mathbf{R}_{\varphi^2} \cdot \mathbf{k}\right)\,.
\end{aligned}
\tag{10}
$$

Formulae (10) are easier to understand if the rotations are interpreted intrinsically. The rate of change of the Euler angles are proportional to the decomposition of torque $\mathbf{M}$ w.r.t. the orientation of the instantaneous reference frame moving/rotating together with the mechanism: a) first the whole mechanism - with its reference frame attached - spins around axis $\mathbf{k}$ then b) around the rotated axis $\mathbf{R}_{\varphi^1} \cdot \mathbf{j}$ and finally c) around the twice-rotated axis $\mathbf{R}_{\varphi^1} \cdot \mathbf{R}_{\varphi^2} \cdot \mathbf{k} = \mathbf{n}$.

For the calculation of $\frac{\partial E}{\partial \varphi^k}$, $k = 1,2,3$ either equations (7) or the equations (10) with torque values (9) integrated over the coordinates $\xi^1, \xi^2, \xi^3$ identified as wand point coordinates by (1) can be used.

**3rd line:** From (2) $\frac{\partial \mathbf{P}_\pm}{\partial \eta^1} = \pm \mathbf{e}_2$ hence we have:

$$I\nabla\left(\mathbf{P}_\pm\right) \cdot \frac{\partial \mathbf{P}_\pm}{\partial \eta^1} = \pm I\nabla \cdot \mathbf{e}_2 \,. \tag{11}$$

**4th line:** With direct calculation, the integrands are given by:

$$\begin{aligned} I\nabla\left(\mathbf{P}_\pm\right) \cdot \frac{\partial \mathbf{P}_\pm}{\partial \alpha^1} &= \left(\pm\zeta\cos\alpha^1 - \rho\sin\alpha^1\sin\phi\right)\left[I\nabla\left(\mathbf{P}_\pm\right)\cdot\mathbf{e}_2\right] \\ &\quad - \left(\zeta\sin\alpha^1 \pm \rho\cos\alpha^1\sin\phi\right)\left[I\nabla\left(\mathbf{P}_\pm\right)\cdot\mathbf{n}\right] \,. \end{aligned} \tag{12}$$

Result (12) can also be viewed as the action of the pseudovectors

$$\mathbf{m}_\pm = \left[\left(\xi^2 \mp \eta^1\right)\mathbf{e}_2 + \xi^3\mathbf{n}\right] \times I\nabla\left(\mathbf{P}_\pm\right) \tag{13}$$

w.r.t. their momentary rotational axes: $-\mathbf{e}_1$ at local coordinates $(0, \eta^1, \xi^3)$ and $\mathbf{e}_1$ at local coordinates $(0, -\eta^1, \xi^3)$ respectively (in the former case, a minus sign is required to satisfy the right-hand rule). Substituting (1) to torque expressions (13) we have

$$\begin{aligned} -\mathbf{m}_+ \cdot \mathbf{e}_1 &= \left[I\nabla\left(\mathbf{P}_+\right)\cdot\mathbf{e}_2\right]\xi^3 - \left[I\nabla\left(\mathbf{P}_+\right)\cdot\mathbf{n}\right]\left(\xi^2 - \eta^1\right) \\ &= \left(\zeta\cos\alpha^1 - \rho\sin\alpha^1\sin\phi\right)\left(I\nabla\cdot\mathbf{e}_2\right) \\ &\quad - \left(\zeta\sin\alpha^1 + \rho\cos\alpha^1\sin\phi\right)\left(I\nabla\cdot\mathbf{n}\right) \\ \mathbf{m}_- \cdot \mathbf{e}_1 &= -\left[I\nabla\left(\mathbf{P}_+\right)\cdot\mathbf{e}_2\right]\xi^3 + \left[I\nabla\left(\mathbf{P}_+\right)\cdot\mathbf{n}\right]\left(\xi^2 - \eta^1\right) \\ &= -\left(\zeta\cos\alpha^1 + \rho\sin\alpha^1\sin\phi\right)\left(I\nabla\cdot\mathbf{e}_2\right) \\ &\quad + \left(-\zeta\sin\alpha^1 + \rho\cos\alpha^1\sin\phi\right)\left(I\nabla\cdot\mathbf{n}\right) \end{aligned} \tag{14}$$

which is equivalent to (12).

### 4 Comparison

The performance of the algorithms was evaluated on 5 acquired image stacks of different pipettes. The pipettes were lowered around the working distance to have similar light conditions to those during the experiments with samples. The pipette tip is moved over at least half of the image and it is assured that the stack always contains it. The voxel position of the tip was manually selected in the stacks. Then the initialization algorithm was run on the stacks and its results were the starting point of PH3D. The results of both the initialization algorithm and PH3D were saved and compared to the manually picked positions. The mean difference between the line profile estimation and the ground truth data is 14.64 ± 4.23 voxels, while for PH3D the error is 8.63 ± 4.80 voxels. The pixel size is 115 nm and the distance between the slices in the stack was 1 $\mu$m, thus the error expressed in micrometers is 0.99 ± 0.55 $\mu$m. This error value is fairly low to reliably hit a cell if the pipette is oriented to its center, since the diameter of cells is usually in the range of 5-20 $\mu$m. Furthermore, this error value is smaller than the reported 3.53 ± 2.47 $\mu$m or 32.97 ± 23.10 pixels in the case of the original 2D model.

### References

[1] Krisztian Koos, József Molnár, and Peter Horvath. Pipette hunter: Patch-clamp pipette detection. In *Scandinavian Conference on Image Analysis*, pages 172–183. Springer, 2017.
