## Supplementary Information: Pressure Regulator for "Automatic deep learning driven label-free image guided patch clamp system for human and rodent in vitro slice physiology"

### Pressure Regulator Setup

*Supplementary Information*

We built a custom pneumatic pressure regulator to apply air pressure on the pipette. The pressure system is equipped with a 50 ml tank in which the requested pressure is set before opening a solenoid valve to the pipette. Two analog pressure sensors (Honeywell, NJ, USA) were used in the system for the closed-loop pressure regulation. One is connected to the tank, the other measures the pressure subsequent to the valve that connects the tank and the pipette. The sensors and the valves are connected to a data acquisition device (DAQ, National Instruments USB-6009, Texas, USA) which controls the valves to set the desired pressure. The pneumatic parts are connected by silicone tubes. Fig. 1 shows the wiring diagram of the system. The electrophysiological signal is also forwarded to the same data acquisition device for real-time detection of the impedance change of the pipette tip.

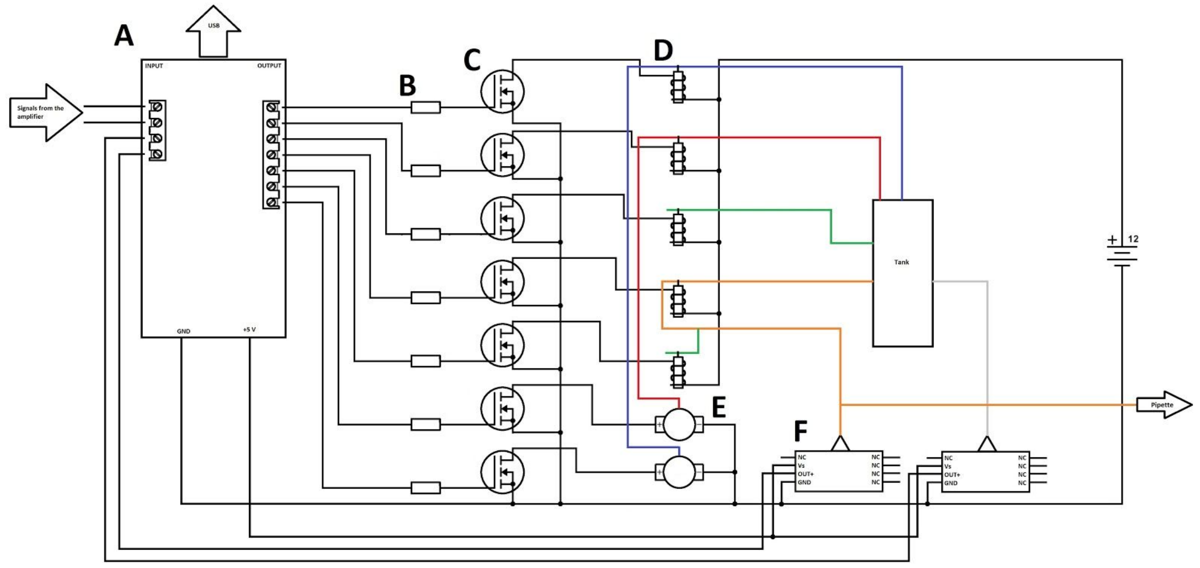


Figure 1: **Wiring diagram of the custom made pneumatic pressure regulator.** A: USB digitizer board (National Instruments, USB-6009), B: 220 Ohm resistors, C: IRFZ44N field effect transistors, D: 12V DC solenoid valves, E: 12V DC pneumatic pumps, F: analog pressure sensors (Honeywell, NJ). Colored lines: pneumatic tubes. red: positive pressure, blue: vacuum, green: atmospheric pressure, orange: desired pressure to the electrode tip, grey: tank pressure. The valves were controlled with TTL signals via the USB board digital outputs. The output voltages of the pressure sensors were measured with the analog input channels of the USB digitizer.

The controller of the pressure system is included in the software. The accuracy of the regulator which is 10 mBar by default can be set from the software. Note that the mixing time can increase if the accuracy is set too high compared to the volume of the installed tank and the silicon tubes. The authors help in providing a kit or an assembled version of the controller on request.
