## Supplementary Information: Software Usage for "Automatic deep learning driven label-free image guided patch clamp system for human and rodent in vitro slice physiology"

### Software usage and parameters

*Supplementary Information*

#### Introduction

This document introduces the autopatcher software to the user and explains the most important parameters which can be set from the graphical user interface. The Installation section describes how to set up the system and install the necessary submodules. Then the subsequent sections demonstrate how to use the different functionalities.

#### Installation

The software is written in Matlab, thus having an installed Matlab R2017a or later is recommended with Image Processing and Computer Vision System toolboxes. Simulator mode works with these licenses, but depending on the hardware setup additional toolboxes or support packages might be required. A webcam needs the support package for USB Webcams, while a FireWire camera requires the Image Acquisition Toolbox and the support package for DCAM 1394 cameras. Our pressure controller system and electrophysiological signal processor uses a National Instruments USB-6009 board which requires Data Acquisition Toolbox and the support package for NI-DAQmx Devices. Initially, the software is configured to run in simulator mode. We provide solutions for the mentioned devices and others can also be applied. If the user has access to some different hardware then developing controllers by inheriting abstract classes makes them available for use with the software. Modification of the existing code is not needed.

The configuration of the software is managed by XML files. The tag values are processed as Matlab code. The configuration supports classes, constructor parameters, most often used data types, and method calls. The main configuration file is vistool_config.xml, but some functionality requires the blindpatcher_config.xml, trainer.xml and prediction_config.xml files. The hardware controllers have to be set up in `<element>` tags before starting the software. Those parameters defined inside the `<properties>` tag are tracked and their most recent values are updated in the file when closing the software.

The autopatcher software requires no other installation than extracting the files to a folder. At this point, the software is ready to be used, however, some functionalities require installation of third-party libraries which are detailed below.

Cell detection requires an installation of Caffe [1]. The provided code uses the Matcaffe interface. Ref. [2] is a guide for installing Matcaffe in newer Matlab versions which is required by our software. Then the Matcaffe folder has to be added to the Matlab path manually. Detection is a computationally expensive task and it is suggested to install Matcaffe on a separate PC with Matlab installation and a CUDA capable GPU card. In this case, the predictor `<element>` in the config file has to be changed from PredictorLocal to PredictorRemote. If the detection is performed in a different Matlab session then startPredictionServer.m has to be run before it accepts tasks. The startPredictionServer script can be configured in prediction_config.xml.

DIC reconstruction requires an OpenCV and OpenCL installation.

### Starting the software: Main window

The software can be started after changing the Matlab working directory to the location of the extracted files. If the startup.m file has not run automatically it has to be run manually. The main window can be started by running the startVisualizationTool.m script which loads the configuration from vistool_config.xml. The logging system might create a log folder and files with .log extension.

Fig. 1 shows the main window in live view mode. The live camera or the loaded image is shown on the left. Around the bottom right area of the image, the z level of the microscope is shown in live mode. On the right side, the most often used buttons can be found. The Live View push-button activates or deactivates the camera. If the Live prediction push button is on the cells are detected in the live camera image. The other buttons are used for visual patch clamping and discussed in the Visual patch clamping section. The menu bar at the top of the window is the starting point for other functions and is discussed element-by-element in the rest of this section.

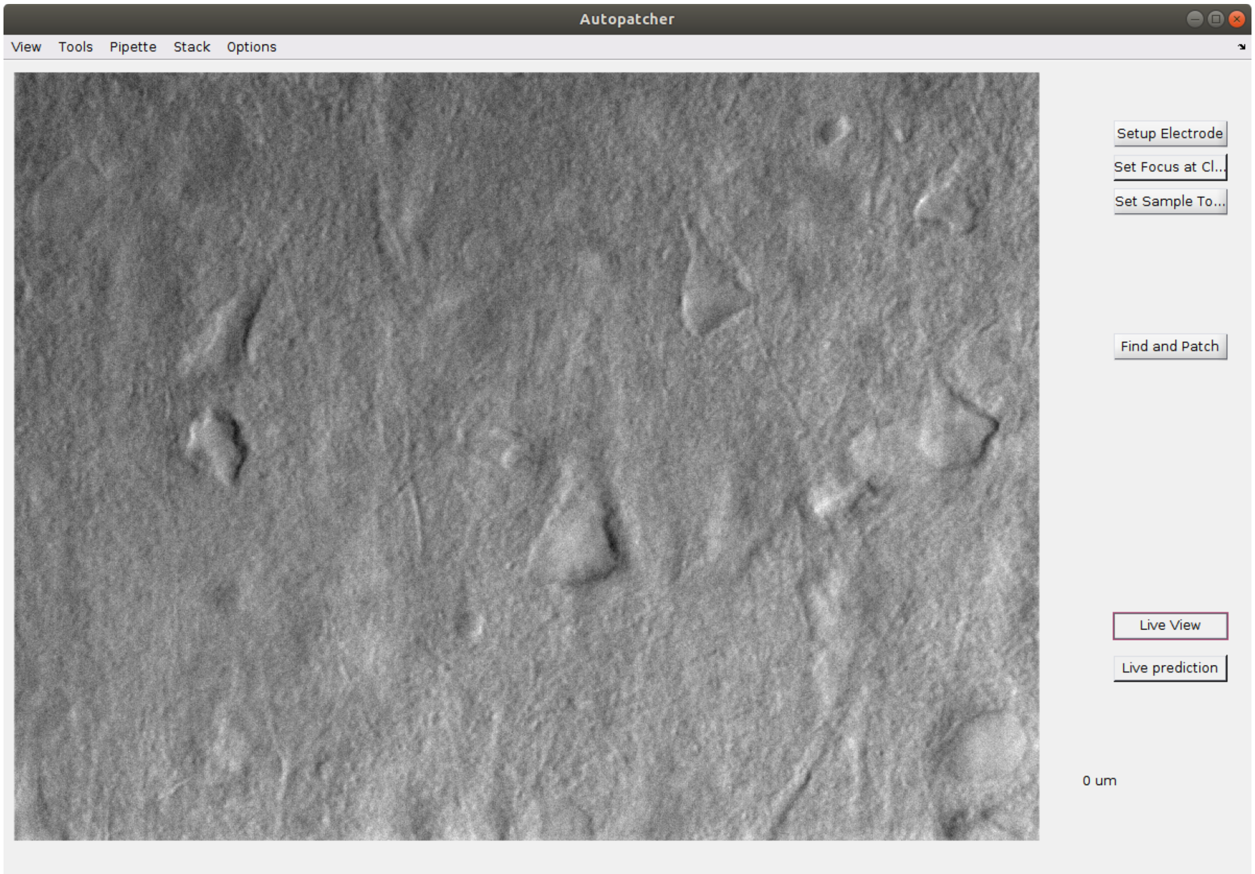


Figure 1: **The main window of the autopatcher software with live view started.**

View menu:

- Blind Patcher: Opens the Blind Patcher window or brings it in focus if already opened.
- Visual Patcher: Opens the Visual Patcher and the Patch Clamp Diary windows, if not opened.
- Cleaner: Opens the Pipette Cleaner window or brings it in focus if already opened.

- Trainer: Opens the Manual3DTrainer window or brings it in focus if already opened and enables its functionality in the main window.

Tools menu:
- Capture and Save image: Saves the current image visible in Live View to a file.
- Reset DCAM: Resets the DCAM driver. Useful if the driver crashes.

Pipette menu:
- Set focus at click: Update the position of the pipette tip by clicking on it in the live camera image.
- Detect focus: Update the tip position of the pipette automatically by a line profile estimation and improve it with the Pipette Hunter 3D 2 wands model.
- Configure: Calibrate the pipette axes for visual patch clamping. This feature should be run after the first startup and before using any visual patch clamping related functionality. A coordinate system transformation has to be determined so that the pipette can be moved based on the microscope's stage coordinate system, or in other words based on the image that we see. This function asks the operator in a step-by-step fashion to move the pipette in every axes at least 100 micrometers (the longer the movements are the more precise the calibration will be) and click on the pipette tip after every movement.
- Ignore sample top: Ignores the sample top position when using the 'Move pipette here' feature in the image and moves the pipette using the 'fast' speed. If the item is unset then the pipette is moved slowly in the tissue and the final step is a forward movement in the x-axis.

Stack menu:
- Load: Loads and shos an image stack.
- Acquire: Acquires and shows an image stack. The number of elements in the stack can be set in the config file under GeneralParameters.stackSize. The current focus level is going to be the topmost image and the step size will be 1 micrometer between the slices.
- Predict: Detects cells in the currently loaded image stack.
- Patch predicted: Offers cells for patch clamping from the recent detection results.
- Show original (default): Shows the original loaded stack (with image normalization).
- Show reconstructed: Performs DIC reconstruction and shows the reconstructed stack.
- Show bg corrected: Performs background correction in the stack and shows the result.

Options menu:
- General options: A window for general options (Fig. 2)
    - Camera timer period: The period in seconds to update the live view screen
    - Acquired stack size: The number of images to acquire in an image stack
    - Save Find&Patch stack: Saves the image stack as a file that is acquired by using the Find&Patch button
    - DIC reconstruction group:
        - Iterations: The number of iterations for the algorithm
        - Direction (degrees): Shear direction in degrees
        - Step size weight: The update step size for the gradient descent method

- Smoothness weight: Multiplier of the smoothness term
- Cores in a thread: The number of CUDA threads in a thread group

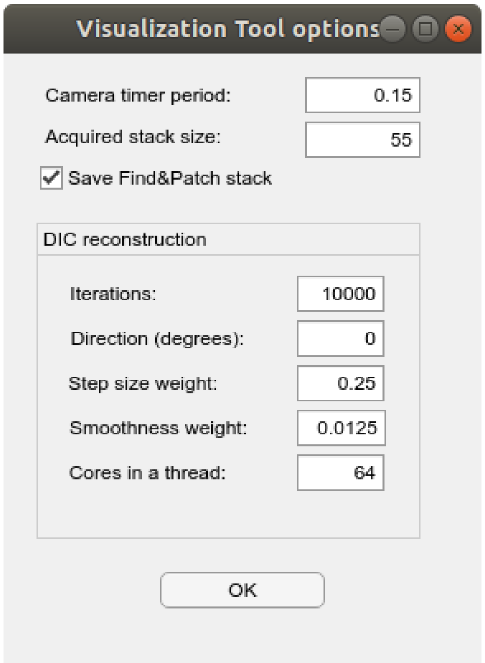


Figure 2: **General options window.**

- Prediction options: Options window for cell detection (Fig. 3)
    - Threshold to detect: The detection result is a probability map that is thresholded with this value to get bounding boxes.
    - Prediction timer: The time interval in seconds to run the detection in live view.
    - Minimum/Maximum object width/height: Detected cells with smaller/bigger bounding boxes are not considered as valid results.
    - Minimum overlap to unite: A ratio, if two detections overlap at least this much in different z slices then they are united.
    - Maximum z distance to unite: If the slices, based on their indices, are further from each other than this value they will not be united.

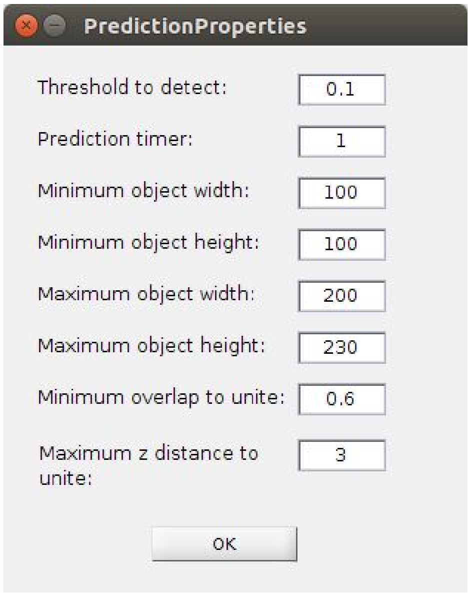


Figure 3: **Prediction options window.**

### Blind patch clamping

Blind patch clamping can be initiated from the Blind Patcher window, which is shown in Fig. 4. The window contains a Pressure panel to check or manually control the pressure system. Before starting any patch clamping it is recommended to check if the pressure and vacuum sources can provide the preset low/high +/- strengths using the related buttons. It is possible that the sensor does not detect perfectly 0 mBar pressure even if it is physically guaranteed. To compensate for this offset (usually between -10 and +10 mBar) run the Calibrate feature which lasts a few seconds, opens the valves to atmosphere, measures, and saves this offset for later calculations. The Off button disables the pressure controller, which is useful if the operator wants to take over control. Beyond the preset pressure values, the operator can request any reasonable strength by entering the number in the Requested pressure field and pressing enter. The pressure measured on the pipette is visible at all times in the Actual pressure field. The status of the pressure controller can be Regulating (normal operation), Calibration, CalibrationComplete, Disabled, BreakIn and BreakInComplete.

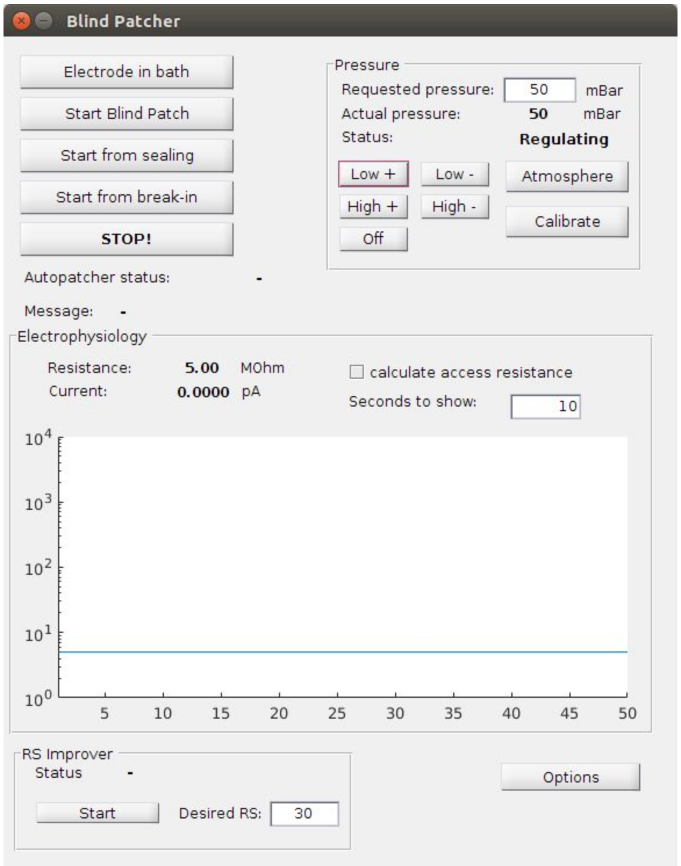


Figure 4: **Blind Patcher window.**

The Electrophysiology panel contains information about the actual resistance and current values and shows the resistance history on a logarithmic scale. Measuring the resistance requires a square command signal which can be activated by the Electrode in bath button in this window.

The Blind Patcher window can be opened in two ways. In both ways it reads the blindpatcher_config.xml file. If it is opened from the Main Window then only the window related properties are used and the control objects are passed to it. It is also possible to perform blind patch clamp recording by starting the startBlindPatcher.m script which opens the window without visual patch clamping capabilities. In this case, the config file has to define the necessary control objects as well.

Performing a blind patch clamp recording starts by clicking on the Electrode in bath button. When pressing this button the pipette should be in the recording solution. This function sets up the pipette by starting a square signal as the command signal for measuring the resistance and applies a small pressure. If the operator finds the resistance value of the pulled glass pipette appropriate, the dura should be crossed by manually moving the pipette and applying higher pressure value. When the pipette is ready for a patch clamp attempt and is near cells the operator has to click the Start Blind Patch button. The current phase of the blind patch clamp process is always visible in the Autopatcher status field in the GUI and can take one of the following three values: Starting, Hunting, Sealing, BreakIn, Success, and Failed. The Message field shows informative details about the current status of the process. In the hunting phase, the pipette is regularly pushed forward while applying a small pressure until the measured resistance value increases. If a relevant resistance increase is detected then the pipette movement is stopped, the pressure is dropped to atmosphere and the phase is changed to Sealing. In the sealing phase, small vacuum is applied to form the gigaseal state between the cell and the pipette. If the phase lasts long, the pressure is set to atmosphere for a few short time intervals, the vacuum strength is increased and the pipette is moved a bit in every direction. When the gigaseal state is achieved the phase is changed to BreakIn. In the break-in phase short but strong vacuum pulses are applied to break the cell membrane and achieve whole-cell configuration. If the break-in was successful the phase is changed to Success and the recording is automatically started. If any of the phases fail, eg. the pipette is moved too much without hitting a cell in the hunting phase, gigaseal could not be formed in the sealing phase, or the cell died during break-in, then the phase is changed to Failed. If the operator finds a problem at any point in the process which was not detected by the system the attempt can be stopped by clicking on the Stop button. If the problem could be fixed the operator can restart the process from any phase with the Start from sealing or the Start from break-in buttons.

The RS Improver panel contains the Start/Stop button and the Desired RS field and tries to improve the series resistance after a patch clamp attempt by alternating between small pressure and small vacuum.

The Options button opens the Blind Patcher Options window shown in Fig. 5. The parameters which can be set here are the following:

- Low positive pressure: Low pressure value used in the patch clamping process.
- High positive pressure: A preset high pressure value (not used in the process but useful for cleaning the pipette tip).
- Low negative pressure: Weak vacuum used in the patch clamping process.
- High negative pressure: Strong vacuum used in the break-in phase.
- ‘Forward’ axis: The axis on which steps are made in the hunting phase
- Hunting group:
    - Min. RS change to seal: Minimum increase in resistance value to switch from hunting to sealing phase
    - Step size: The step size with which the pipette is pushed along its forward axis every second.
    - Maximum distance: The maximum distance the pipette can take in the hunting phase. 0 means no limit.
    - Check hit reproducibility: If checked, pulls back the pipette after a hit; this might increase hit robustness
    - Pull back steps for reproducibility: Number of steps to pull back
    - Clog warning R increase: Warns the user if the R increases by this amount but no hit was detected
- Sealing group:
    - Check R increase on atmosphere: Checks if R increases on atmosphere without applying vacuum, stops otherwise. The amount of increase should be at least as much as ‘Min. RS change to seal’.
- Break-in group:
    - Initial break-in delay: The desired length of first vacuum pulse in the break-in phase.

- Break-in delay increase: The desired increase of the duration of the vacuum pulse after every break-in attempt.
- Gigaseal resistance value: Minimum resistance value which is considered a gigaseal. Measuring higher resistances has higher error rates, thus setting this value higher than the theoretical 1 GOhm is reasonable.
- Successful break-in resistance value: Maximum resistance value which is considered a successful break-in attempt / whole-cell patch.
- Minimum delay before first break-in: Delay between a formed gigaseal state and the first break-in pulse attempt.
- Break-in phase total length: The total length of the break-in phase to allow break-in attempts.
- Pull back after attempts: Pulls back the pipette after this number of break-in attempts. It helps if the nucleus is at the pipette tip and it prevents the formation of a gigaseal.
- Pull back distance: The desired extent  the pipette should be pulled back after a few break-in attempts.

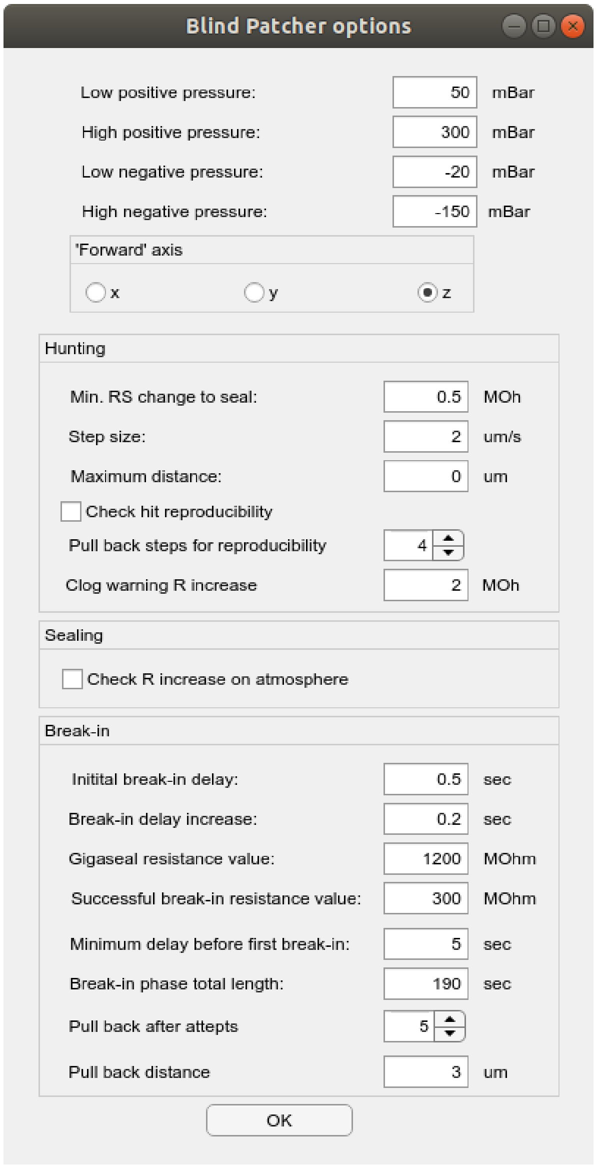


Figure 5: **The Blind Patcher options window.**

### Visual patch clamping

Visual patch clamping is the process of visually selecting a cell and performing recording on it. The system automatically tracks the shift of the cell in the tissue due to the deformation caused by pushing the pipette deeper, and avoids obstacles, like blood vessels and other cells, with the pipette on its way to the target.

The most often used way to start visual patch clamp recording is through the buttons on the Main window. It is assumed that the operator has already configured the pipette and moved it in the recording solution close to but above the sample. First, the operator has to click on the Setup Electrode button which applies pressure and starts the square signal. In practice, it has the same functionality as the Electrode in bath button in the Blind Patcher window. Then the pipette tip position has to be updated (especially if a new pipette has been inserted). The more consistent way of updating the tip position is running the Pipette Hunter algorithm from the Pipette menu -

Detect focus item. However, depending on the host machine, for an experienced operator it might be a few seconds faster to click on the Set Focus at Click button and then click on the tip in the camera image. Then the focus level should be set to the sample's top and the Set Sample Top Here button should be clicked to save this position. Finally, the Find and Patch button has to be pushed to run the cell detection and offer cells for recording. The detected cells are offered in descending order of the detection certainty. Fig. 6 shows the Main window when the Find and Patch button is pushed and the system offers cells for recording for the operator. The selected cell is highlighted with a dashed bounding box.

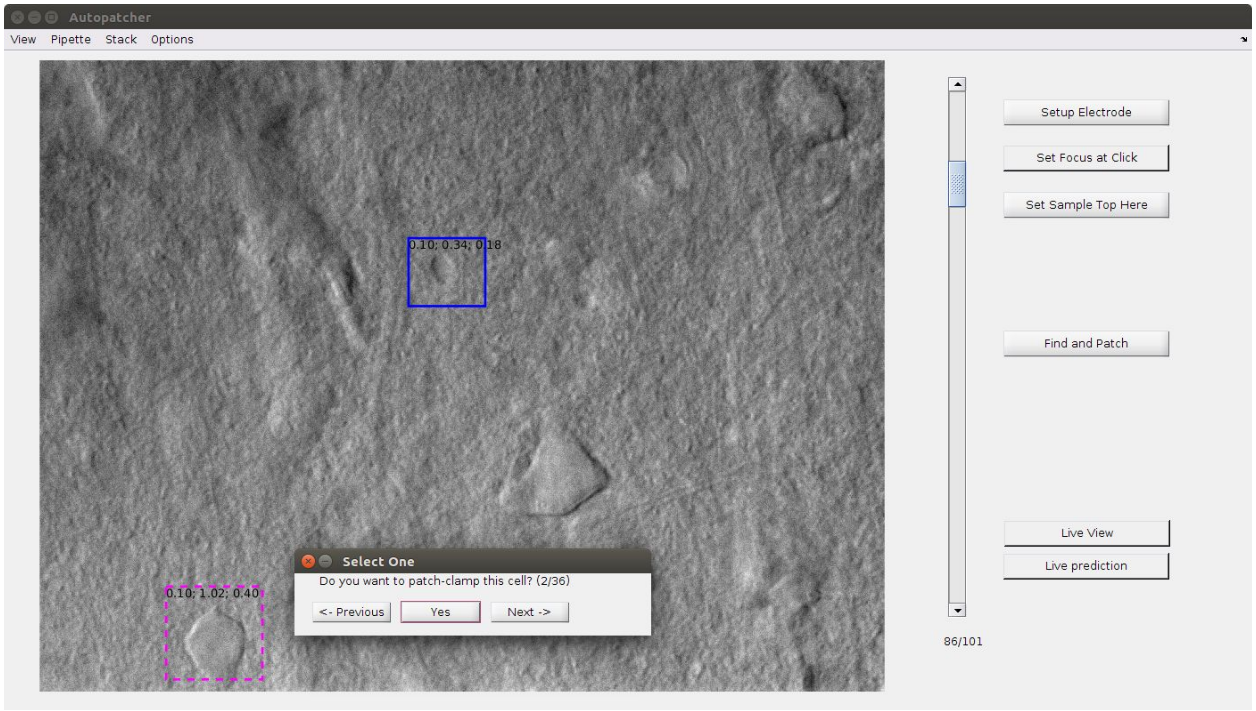


Figure 6: **Using the Find and Patch feature.**

The other way of starting visual patch clamping is to right-click on a cell in the camera image and select the 'Do patch-clamp here' item from the context menu as shown in Fig. 7. This approach does not detect cells and the operator should aim to the centroid of the cell to maximize the chance of the pipette hitting the cell.

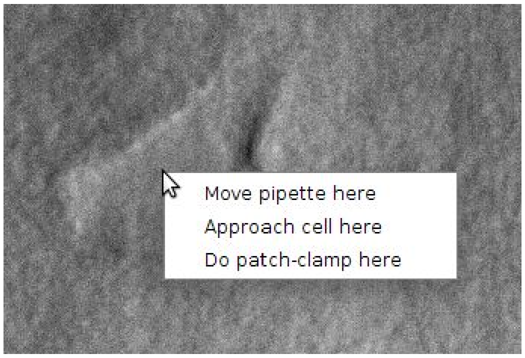


Figure 7: **Starting visual patch clamp recording without cell detection.**

After the visual patch clamping is launched, multiple subsystems are started. The selected cell is centered in the camera image, the pipette is oriented so that only a forward movement in its X axis is needed to approach the cell. The cell tracking system draws a red square around the cell to note which area around the cell is tracked. A middle pressure is applied and the pipette is regularly moved closer to the cell. Meanwhile, the obstacle avoidance system helps to dodge objects on the way to the target cell. The Visual Patcher Control and Blind Patcher windows are opened or brought into focus for the operator to supervise the process. The Visual Patcher Control window shown in Fig. 8 contains information related to the mentioned subsystems. When the pipette is just a few micrometers from the target, the Visual Patcher stops and starts the Blind Patcher which performs patch clamp recording. We found that the cell often shifts if the blind patch clamping is performed on the X axis of the pipette. Therefore the system is configured so that the approach places the pipette above the cell and the blind patch clamping is performed by pushing the pipette on the cell from above on the Z axis. However, this behavior can be configured in the configuration file.

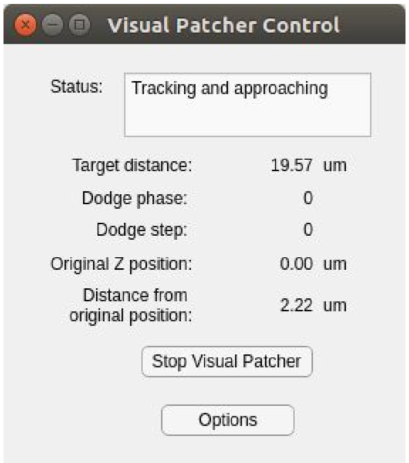


Figure 8: **The Visual Patcher Control window.**

The Options button in the control window opens the Visual Patcher Options window which is shown in Fig. 9. The adjustable parameters are detailed below.

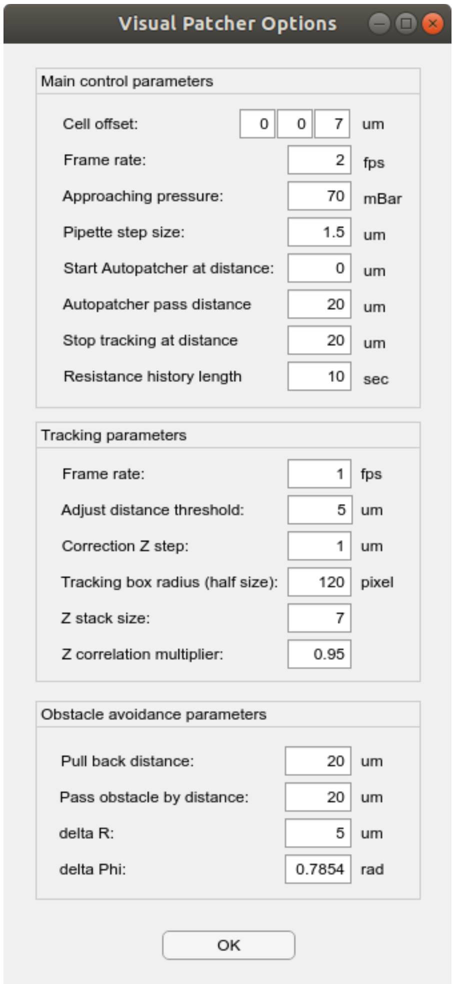

Figure 9: **The Visual Patcher Options window.**

Main control parameters:

- Cell offset: An offset value can be set on all three axes where the pipette will be oriented, relative to the actual position of the cell. This parameter is useful when the forward axis during blind patch clamping is ‘z’ thus a ‘z’ offset should be set a few micrometers above the cell.
- Frame rate: The rate per second when the control function should make a step by pushing the pipette forward, look for obstacles and check the remaining distance to the target.
- Approaching pressure: The pressure value used when approaching the cell. Usually, it is higher than the low positive pressure used for blind patch clamping, since the target can be approached faster.
- Pipette step size: The distance the pipette should be pushed forward when the control callback function is called.
- Start Autopatcher at distance: The distance between the pipette tip and the target cell when blind patch clamping should be started. The control function stops pushing the pipette at this distance. Usually, the value is set in combination with Cell offset. If the blind patcher’s forward axis is ‘z’ and the Autopatcher is started at 0 distance, a cell offset in Z axis of 5-7 micrometers should be used.

- Autopatcher pass distance: The maximum distance the Autopatcher is allowed to take once it is started and before it is stopped by the controller function. 0 means no limit.
- Stop tracking at distance: The distance between the pipette and the target when tracking should be stopped. If the pipette is close to the cell it usually affects the image around it which has negative effects on the tracking system. Usually, this value is set to the same as the Tracking box radius value in the Tracking parameters.
- Resistance history length: The length of the resistance value history, in seconds, that should be kept. This vector is used, for example, to detect obstacles.

Tracking parameters:

- Frame rate: The rate per second the tracking system should make a step at. The tracking steps alternate between xy (lateral) tracking, taking a small image stack, and z tracking in the stack.
- Adjust distance threshold: The tracking system does not correct every little shift of the tracked object but saves this information. The system only moves the pipette when the shift of the target from its original position to the actual one, in micrometers, is more than this value.
- Correction Z step: The tracking system in the Z axis can only indicate whether the target shifted up or down. The correction step of the pipette will be this value if a shift in the Z axis is detected.
- Tracking box radius (half size): Half size of the box in pixels which should be used for tracking in the image. It should be about as big as most cells in the image (thus the box will be twice as big). Depending on the Pipette step size parameter the value can be increased or decreased.
- Z stack size: The number of slices in the z stack which is acquired for tracking in the Z axis.
- Z correlation multiplier: The tracking system uses correlation to determine the shift in the Z axis. However, if the image is noisy the correlation value will also be affected. This multiplier is applied to the correlation value of image slices other than the middle and shift is only detected if they are still higher than the correlation of the middle and the reference images.

Obstacle avoidance parameters:

- Pull back distance: If an obstacle is detected the pipette is pulled back to the extent of this value in micrometers.
- Pass obstacle by distance: An obstacle avoidance attempt is finished when the pipette passes the previously detected obstacle by the distance given in this parameter. Note that this value is not related to the Pull back distance parameter; after the pipette is pulled back it will be pushed forward by both the pull back and pass distances together.
- Delta R: A distance factor used in the formula described with Delta Phi for lateral movement along the axis where the pipette moves during obstacle avoidance [3].
- Delta Phi: A rotation factor used for the formula below for lateral movement along the axis where the pipette moves during obstacle avoidance:

$$m(n) = n\Delta r \cos(n\Delta\Phi - \pi/4)\mathbf{i} + n\Delta r \sin(n\Delta\Phi - \pi/4)\mathbf{j}$$

### Patch Clamp Diary

The Patch Clamp Diary subsystem automatically collects information during the use of the software for patch clamping and is able to visually show the attempt positions and generate statistics files. The system intends to replace a paper-based laboratory notebook. Besides the positions, the system saves the outcome and timestamps of the phases of the attempts. The system can be used with or without the visual patch clamping functionalities.
The Patch Clamp Diary window can be opened along with the Visual Patcher window from the main window using the View - Visual Patcher item. The window has three tabs: Slice info, JEM slice info, and Patch-Seq Pipette. Fig. 10 shows the window with the Slice info tab activated which enables the marking of the sample's side and holder positions and displays them along with the positions of patch clamp attempts.

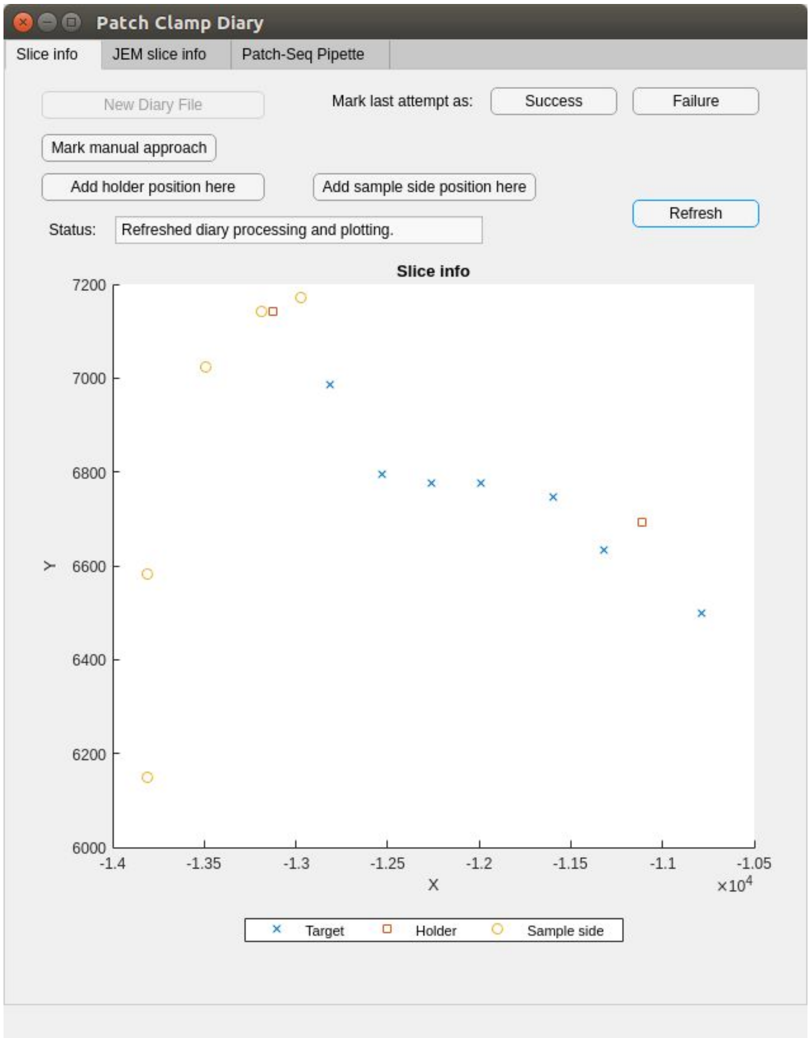


Figure 10: **Patch Clamp Diary window when the Slice info tab is active.**

The Slice info drawing shows the saved positions and has a figure legend. The Status field shows useful information about the function being used. For example, when a holder position is added the Status field outputs the saved position or shows an error message if the position could not be queried but does not interrupt the work of the operator. The role of the buttons and elements in the Slice info tab is detailed here:

- New Diary File: Generates statistics and saves it to a CSV file. It also makes a backup file of the raw log data with the same name but a different extension than the statistics file. Note that this button should be used at the end of a patch clamping session, when finished with a sample or replacing it with a new one, to generate statistics of the last used.
- Mark last attempt as Success/Failure: The outcome of an attempt can be set manually using these buttons. This is useful if the system did not detect an early break-in or if the operator performed patch clamping manually.
- Mark manual approach: If the operator does not use the system's visual patch clamping functionality (blind patcher only or manual) the position of the new attempt can be set using this button. The saved positions are always in the stage coordinate system and the target cell should be approximately in the center of the image.
- Add holder position here: Saves the current stage position as a holder position. It is a good practice to place the holder approximately in the center of the image before clicking on this button.
- Add sample side position here: Saves the current stage position as a sample side position. It is a good practice to place the side of the sample approximately in the center of the image before clicking on this button.
- Refresh: Generates statistics, saves it to the default file name and updates the Slice info drawing.

The generated CSV file contains column names, most of which are self-explanatory but some of them are explained here. The TargetDepth value is measured from the manually set sample top position, while the TargetDistance expresses the distance the pipette should move on its X axis to reach the target. The abbreviation 'AP' in multiple columns stands for Autopatcher which is the blind patcher subsystem. The DetectionSelectedIndex shows the index of the selected cell when using the Find and Patch function. The timestamps are in the host system's timezone. Positions and distances are expressed in micrometers, resistance values are in megaohms.

The JEM Slice info and Patch-Seq pipette tabs are used together to document nucleus harvesting attempts. Fig. 11 shows these tabs. Most values should be filled manually but there are some which are filled automatically, e.g. timestamps and resistance values.

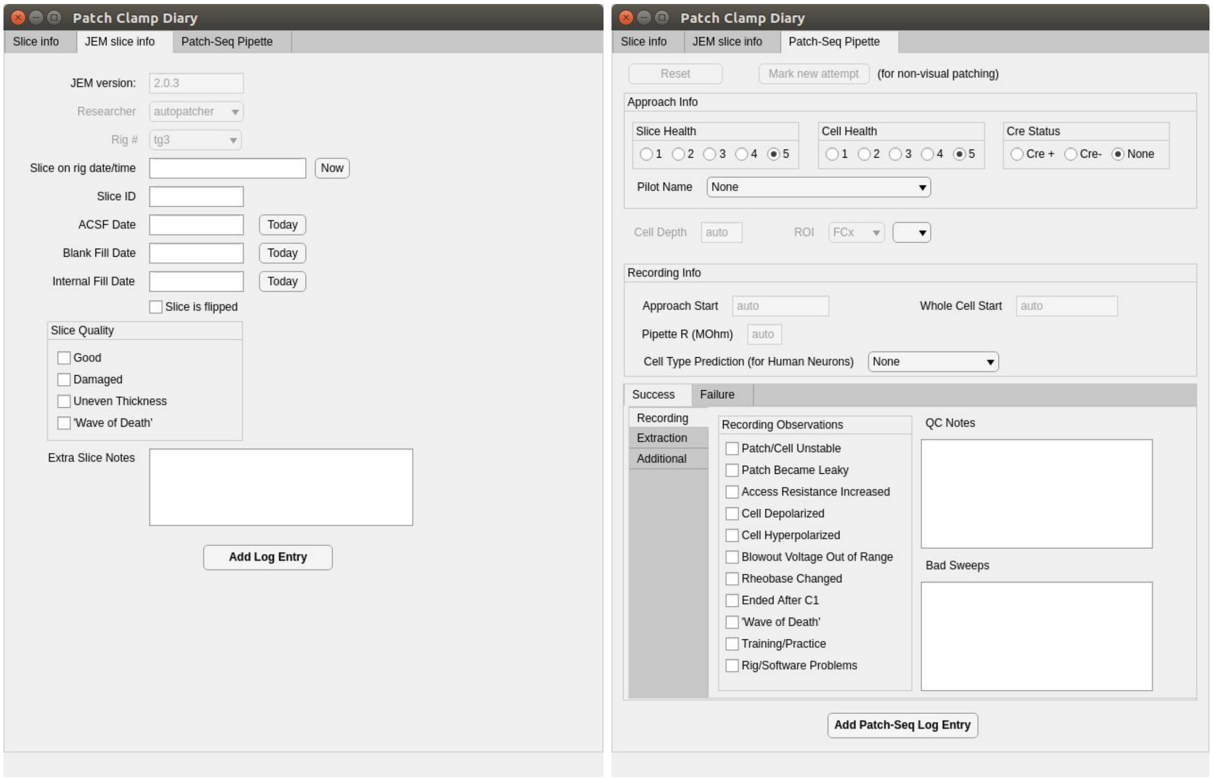


Figure 11: **The JEM slice info and Patch-Seq Pipette tabs.**

### Pipette Cleaner

The pipette cleaning functionality [4] can be configured and used from the Pipette Cleaner window which can be opened from the main window's View - Cleaner item and is shown in Fig. 12.

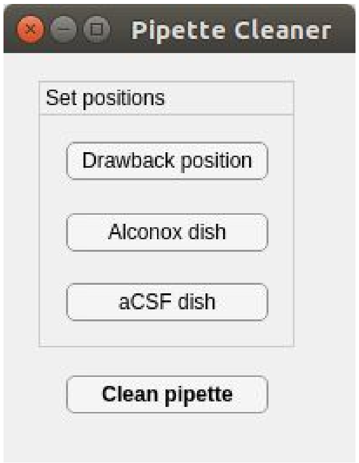


Figure 12: **The Pipette Cleaner window.**

Before cleaning a pipette with the functionality the cleaning detergents have to be installed around the sample. Two detergents are required: standard recording solution (aCSF) and

Alconox. We have designed and 3D printed an object, shown in Fig. 13, that can hold two 250 µl PCR tubes containing the detergents and can be attached to the objective. Other tube holders or even Petri dishes can be used.

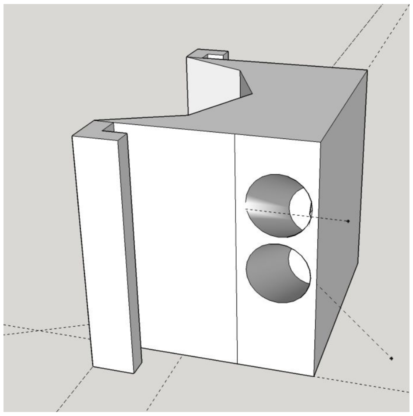

Figure 13: **Schematic of the 3D printed object that holds the cleaning detergents.**

When the detergents are installed and the objective is lowered to the position where a sample would be visible, the operator should make sure that the pipette can reach the tubes and make adjustments if necessary. Then the pipette has to be calibrated for the dish positions using the GUI which are saved and loaded the next time the software starts. The Drawback position button saves the X position of the pipette which should be set such that the objective or other objects are not hit around the tubes when moving the pipette in the other axes. Then the pipette should be manually moved to the tube containing Alconox such that the tip is immersed with the detergent. The Alconox dish position button saves the position of the pipette. This has to be repeated for the other liquid before clicking on the aCSF dish button. Then the pipette can be cleaned after patch clamp recording attempts by the Clean pipette button. This button starts the process by pulling back the pipette to the drawback position, adjusting it laterally before pushing it in the dishes and alternates between blowing and suction pulses to clean the pipette. When the cleaning is finished the pipette is moved back safely to the position where the process was initiated.

### Image stack labeling tool

We developed a tool for image stack labeling in the main window of the software. The tool allows the user to put 2D bounding boxes around the cells in the images over multiple stacks. We used the labels generated with this tool to teach a convolutional neural network for DIC cell detection. The tool can be started from the main window's View - Trainer item and Fig. 14 shows it while a labeling is in progress.

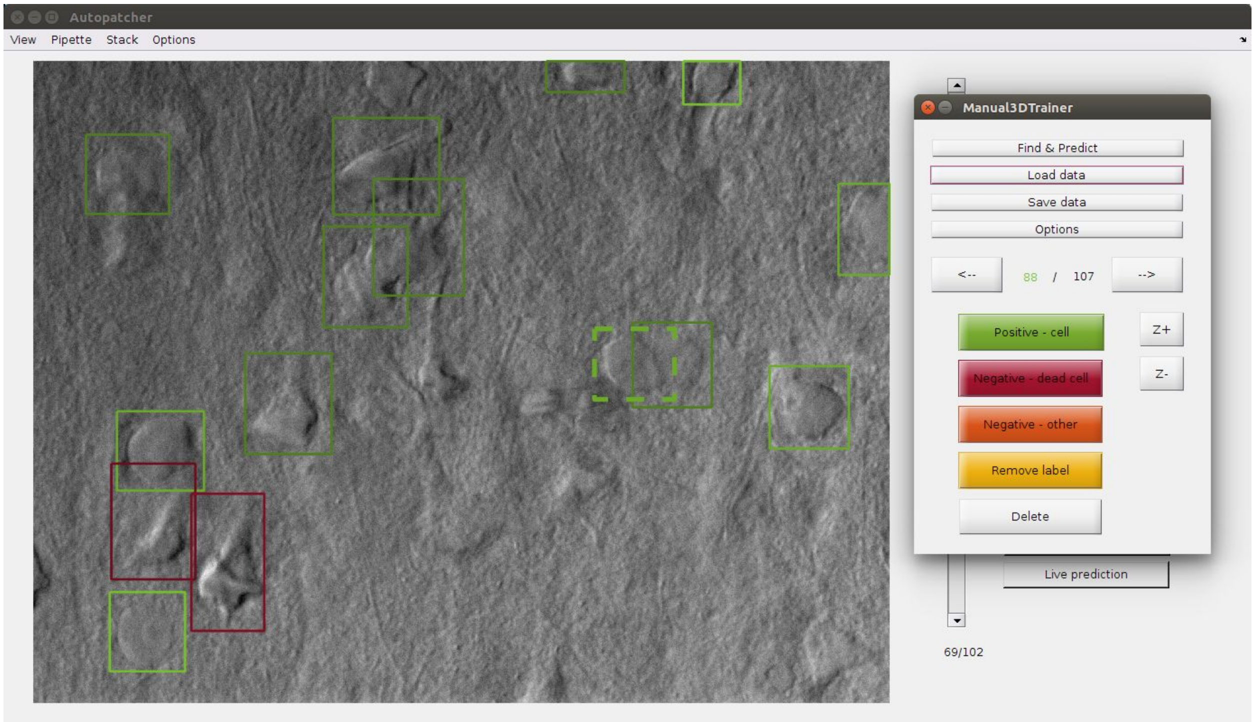


Figure 14: **The image stack labeling tool.**

The tool can only be opened if an image stack is loaded (or acquired) in the main window. When the tool is opened, the labeling functionalities are enabled. Double-clicking in the image adds a new label with a default box size. The dashed lines always indicate the selected box. The edges of the boxes can be adjusted by dragging them with the mouse while the control key is pushed. The extent of the boxes in the Z dimension can be changed globally but not individually. Thus the boxes should be added in the slice where the cell is in focus. However, the Z position of the boxes can be adjusted later. The boxes can have different labels: a positive for cells, a negative for dead cells, and another negative possibly for other objects like red blood cells. A label can also be removed while the box is not deleted which is useful if the user wants to keep it until a later decision. The data can be saved and loaded and if the cell detection system which uses a previous network is installed it can be used to help the user and automatically put bounding boxes on the detected cells. The default new box size and the global size in Z dimension can be set and the values are saved in the trainer.xml file in the config folder.

Below are the description of the buttons in the window:

- Find cells: Uses the cell detection system and puts bounding boxes on the detection results.
- Load data: Loads a previously saved data of bounding boxes.
- Save data: Saves the bounding boxes from the current session to file.
- Options: Opens a dialog where the default box sizes can be set.
- Arrow buttons: Select the next/previous bounding box (dashed line shows the one selected). Arrow keys also switch the selection.
- Z+/Z-: Adjusts the Z position of the selected box.
- Positive - cell: Labels the selected box as positive.
- Negative - dead cell: Labels the selected box as a dead cell.

- Negative - other: Labels the selected box as other negative example.
- Remove label: Removes the label from the selected box but keeps the box itself.
- Delete: Deletes the selected box and its label. The delete key also has this functionality.

### References

[1] http://caffe.berkeleyvision.org/
[2] http://www.cs.jhu.edu/~cxliu/2016/compiling-matcaffe-on-ubuntu-1604.html
[3] Stoy, William Andrew, et al. "Robotic navigation to subcortical neural tissue for intracellular electrophysiology in vivo." Journal of neurophysiology 118.2 (2017): 1141-1150.
[4] Kolb, I., et al. "Cleaning patch-clamp pipettes for immediate reuse." Scientific reports 6 (2016): 35001.
